# An RFX transcription factor regulated ciliogenesis in the progenitors of choanoflagellates and animals

**DOI:** 10.1101/2022.11.11.515474

**Authors:** Maxwell C. Coyle, Adia M. Tajima, Fredrick Leon, Semil P. Choksi, Ally Yang, Sarah Espinoza, Timothy R. Hughes, Jeremy F. Reiter, David S. Booth, Nicole King

## Abstract

Little is known about the origins of the transcriptional modules that coordinate cell-type specific functions in animals. The controlled expression of one cellular feature – the cilium – was likely critical during early animal evolution. Two key transcription factors, RFX and FoxJ1, coordinate ciliogenesis in animals but are absent from the genomes of most other ciliated eukaryotes, raising the question of how the transcriptional regulation of ciliogenesis has evolved. To reconstruct the evolution of the RFX/FoxJ1 transcriptional module and its role in the regulation of ciliogenesis, we investigated RFX and FoxJ1 function in one of the closest living relatives of animals, the choanoflagellate *Salpingoeca rosetta*. Targeted disruption of the *S. rosetta* RFX homolog *cRFXa* resulted in delayed cell proliferation and aberrant ciliogenesis, marked by the collapse and resorption of nascent cilia. Ciliogenesis genes and *foxJ1* were significantly down-regulated in *cRFXa* mutants, consistent with a pre-animal ancestry for this transcriptional module. We also found that cRFXa protein preferentially binds to a sequence motif that is enriched in the promoters of *S. rosetta* ciliary genes and matches the sequence motif bound by animal RFX proteins. These findings suggest that RFX coordinated ciliogenesis before the divergence of animals and choanoflagellates, and that the deployment of this module may have provided a mechanism to differentiate ciliated and non-ciliated cell types in early animal evolution.

## Introduction

Cilia provided our protistan ancestors with the ability to sense and explore environments, avoid predation, and capture bacterial prey^1–3^. However, ciliary motility is energetically costly^4^ and ciliary development can compete with other physiological processes, including cell division and amoeboid motility^2, 5, 6^. In modern animals, deploying cilia in the right cells at the right developmental time point is crucial. In addition, cell biologists have long postulated that the distinction between ciliated somatic cells and non-ciliated germ cells was the first cell differentiation event in the earliest animals^2, 6–9^. Therefore, reconstructing the regulation of ciliogenesis in the ancestors of animals can help to trace the roots of cilia regulation in modern animals, and perhaps the evolutionary foundations of animal cell differentiation.

Despite the lack of a robust fossil record for animal origins, we can infer key features of the progenitors of animals by comparing their biology with that of their closest relatives, the choanoflagellates^5, 10–13^. Choanoflagellates are microeukaryotes with both unicellular and colonial forms, as seen in the emerging model *Salpingoeca rosetta*^14–17^. Choanoflagellate cells feature a distinctive “collar complex” composed of a single apical flagellum surrounded by a collar of actin-filled microvilli^10, 12^. (The terms “cilia” and “flagella” are used synonymously; while “flagella” is more commonly used in the choanoflagellate field, we hereafter use derivatives of “cilia” to align with studies of ciliogenesis in other organisms.) Although choanoflagellates exist most frequently in a ciliated state, they can reversibly transition to a non-ciliated amoeboid form in response to confinement^5^. Additionally, the cell cycle itself requires regulated ciliogenesis in choanoflagellates, ciliated cells in animals, and many other ciliated eukaryotes, as a single pair of basal bodies nucleates microtubules for both the cilium and the mitotic spindle^6, 18^, requiring cells to retract their cilium prior to cell division and re-construct the cilium after each round of mitosis^12, 15, 18^.

The structural components of cilia are remarkably conserved. Eukaryotic cilia feature an axonemal array of doublet microtubules, assembly by intraflagellar transport complexes, and dynein-driven motility^18–21^, all built from dozens of highly conserved genes. These molecular commonalities suggest that the last common ancestor of eukaryotes had a cilium^1, 22^ and that the cilia of choanoflagellates and animals are homologous^23^.

Despite the structural homology of eukaryotic cilia, the key transcription factors (TFs) that regulate animal ciliogenesis – RFX and FoxJ1 – are either missing (e.g. *Chlamydomonas, Naegleria*, ciliates), of unknown function (choanoflagellates, chytrids, amoebozoa), or of non-ciliary function (ascomycete fungi) outside of animals (Fig 1A). RFX TFs regulate the development of both motile and non-motile cilia throughout bilaterian diversity^24–42^, with loss of RFX function affecting the transcription of dozens to hundreds of ciliary genes^31, 34, 41^ and resulting in truncated, malformed, or missing cilia^24, 28, 32, 35, 36, 40^. The up-regulation of RFX TFs in ciliated cells of ctenophores and cnidarians^43, 44^ suggests that RFX may also regulate ciliogenesis in early branching animals. In ascomycete fungi, which do not have cilia, RFX controls cell cycle progression^45, 46^ and DNA damage response checkpoints^47–49^.

**Figure 1.**
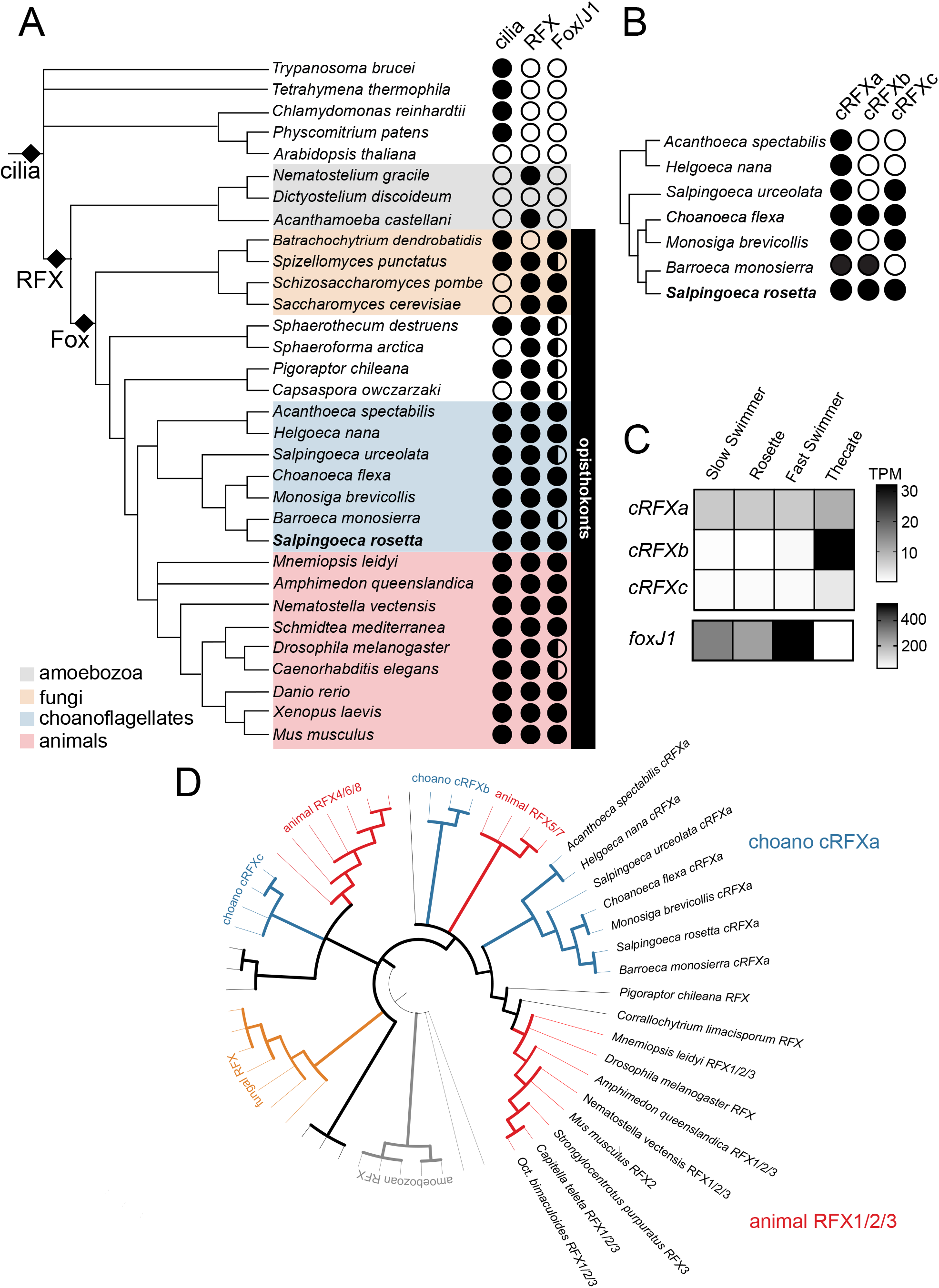
The evolutionary history of cilia-associated transcription factors and their expression in choanoflagellates. (A) Cilia evolved before the emergence of RFX and FoxJ1 TFs. The presence (filled circle) or absence (open circle) of RFX and Fox domain proteins is indicated for diverse eukaryotes (Supp Files 1, 2, Materials and Methods). Cilia are found in most eukaryotic lineages, indicating a cilium was present in the last eukaryotic common ancestor. RFX TFs are found across opisthokonts and amoebozoans and are ubiquitous in choanoflagellates, while Fox TFs are mostly restricted to opisthokonts. See Supp Note 2 for rare exceptions to these patterns. Half shading in the Fox/J1 column indicates the presence of Fox family members, while full shading indicates the presence of a reciprocal best BLAST hit with either the *Xenopus laevis* or *Schmidtea mediterranea* FoxJ1 (Supp File 2, Materials and Methods). Likely FoxJ1 orthologs are detected in most choanoflagellate species. Species tree built from broad eukaryotic and clade-specific phylogenies^124–128.^ ^(B)^ The *cRFXa* sub-family is widespread in choanoflagellates. RFX family relationships were determined using maximum-likelihood phylogenetic trees built by IQTREE^114^ (Supp Fig 2, Supp File 1). All RFX TFs in choanoflagellates grouped into one of three well-supported sub-families: *cRFXa*, *cRFXb*, and *cRFXc.* For representative choanoflagellates, the presence (filled circle) or absence (open circle) of each sub-family are indicated. While *cRFXa* was detected in all cultured choanoflagellates that have been sequenced (Supp Note 1), *cRFXb* and *cRFXc* were restricted to subsets of choanoflagellate diversity. (C) *cRFXa* is expressed in all surveyed *S. rosetta* life history stages. *S. rosetta* can transition between multiple colonial and solitary cell types^15^. All stages depicted here bear motile cilia. RNA sequencing shows that of the RFX TFs, only *cRFXa* is expressed above background levels (average TPM [transcripts per million] >= 1) in all cell types. *cRFXb* and *cRFXc* are only expressed above background levels in thecate cells (Supp File 3). *foxJ1* is expressed in all cell types and is highest in fast swimmers (Supp File 3). Shading indicates average TPM value of the gene across three biological replicates. Note the separate scale bars for *RFX* and *foxJ1* expression levels due to the approximately ten-fold difference in maximum expression level between these genes. (D) Choanoflagellate *cRFXa* genes form a clade with the animal *RFX1/2/3* family. This maximum-likelihood phylogenetic tree includes RFX sequences from diverse opisthokonts and amoebozoans (Materials and Methods, Supp File 1). Width of branches indicates bootstrap support and all nodes with less than 75% bootstrap support are collapsed. The clade uniting choanoflagellate *cRFXa* and animal *RFX1/2/3* has 79% bootstrap support. See Supp Fig 3 for full annotated version of this phylogeny.

Despite the different processes regulated by animal and fungal RFX, the DNA-contacting residues of RFX DNA-binding domains (DBDs) are highly invariant^50–52^ (Supp Fig 1), and therefore the RFX monomeric recognition sequence – GTTRCY – is conserved across fungi and animals^53–56^. RFX binding sites often occur as tandem inverted repeats, forming a palindromic sequence referred to as an “X-box”^38, 41, 50, 54, 57, 58^, which is bound by a dimer of RFX TFs^51, 53^.

The second well-studied transcriptional regulator of animal ciliogenesis is the forkhead box J1 (FoxJ1) transcription factor^59^. Depletion or knockout of FoxJ1 in mice^60–62^, frogs^63, 64^, and zebrafish^35, 63^ results in widespread loss of motile cilia and severe developmental defects. This connection between FoxJ1 and ciliogenesis may be conserved across bilaterians; RNAi of the *foxJ1* ortholog in the planarian *Schmidtea mediterranea* blocks the development of motile cilia^65^. Strikingly, ectopic FoxJ1 in zebrafish is sufficient to convert primary cilia into motile cilia^35, 66^.

Homologs of RFX and FoxJ1, two canonical transcriptional regulators of ciliogenesis in animals, have previously been found in non-animals, including in choanoflagellates^47, 51, 52, 67–70^. RFX has been detected across opisthokont and amoebozoan diversity^51, 52^, while Fox genes appear to be restricted to opisthokonts^67–69^ (Fig 1A, Supp Note 1).

Although choanoflagellates express RFX and FoxJ1, the role of these TFs in choanoflagellates has been unclear. One prior study looked for X-box sequences in the promoters of a handful of ciliary genes in *Monosiga brevicollis*, the only choanoflagellate with genome-scale data available at the time^52^. Two out of the twelve assayed ciliary gene promoters in *M. brevicollis* had X-box consensus sequences, but this was interpreted as background signal and the limited available data suggested that RFX gained control of ciliary genes in animals after their divergence from choanoflagellates^52^.

Because animal ciliogenesis regulators are conserved in choanoflagellates, we investigated to what extent this conservation extends beyond the gene level to the gene regulatory networks involved in ciliogenesis, using the genetically tractable choanoflagellate *S. rosetta*^71, 72^. Here we show that targeted disruption of an RFX homolog in *S. rosetta* results in aberrant ciliogenesis and widespread down-regulation of conserved ciliogenesis genes, including *foxJ1*. Moreover, we found that the conserved RFX binding motif is enriched in the promoters of *S. rosetta* ciliome genes. The conserved function of RFX as a regulator of ciliogenesis in choanoflagellates and animals suggests that the RFX regulatory module predates animal origins.

## Results

### Choanoflagellates express orthologs of animal cilia-associated transcription factors

By searching the EukProt^73^ database, an updated and taxonomically rich database that includes all currently available choanoflagellate genomes and transcriptomes, we gained a more complete picture of FoxJ1 and RFX evolution in choanoflagellates (Fig 1A, Supp Files 1,2). Previous work identified an ortholog of FoxJ1 in *S. rosetta*^69, 70^. We confirmed that this gene is a likely ortholog of animal FoxJ1s (as a reciprocal best BLAST hit) and we identified candidate FoxJ1 orthologs in diverse other choanoflagellates (Fig 1A, Supp File 2).

We found that nearly all choanoflagellate express at least one RFX homolog, and many express up to three RFX homologs (Fig 1B, Supp File 1). To determine the relationships among the various choanoflagellate RFX genes, we performed maximum-likelihood phylogenetic analyses. All choanoflagellate RFX genes fell into one of three paralogous sub-families, provisionally named *cRFXa*, *cRFXb*, and *cRFXc* (Fig 1B, Supp Fig 2). Only *cRFXa* homologs were detected in the predicted proteomes of nearly all choanoflagellate species analyzed (see Supp Note 2 for exceptions), while *cRFXb* and *cRFXc* homologs had more restricted phylogenetic distributions within choanoflagellates (Fig 1B). At least eight species, including *S. rosetta,* were found to express representatives of all three choanoflagellate RFX sub-families (Fig 1B, Supp Fig 2, Supp File 1).

The life history of *S. rosetta* includes transitions between diverse cell types – including benthic “thecate” cells, slow swimmers, fast swimmers, and multicellular rosettes^15^ – all of which are ciliated. We found that *cRFXa* was transcribed in each of these life history stages, while *cRFXb* and *cRFXc* expression was restricted to thecate cells (Fig 1C, Supp File 3). Interestingly, *foxJ1* was down-regulated in thecate cells and up-regulated in fast swimmers, a cell type with longer cilia, a faster swimming velocity, and the capacity for pH-taxis^74, 75^ (Fig 1C).

To determine the relationships between choanoflagellate and non-choanoflagellate RFX sub-families, we included diverse opisthokont and amoebozoan RFX sequences in a second round of phylogenetic analysis. Fungal and amoebozoan RFX genes clustered separately from animal and choanoflagellate RFXs (Fig 1D, Supp Fig 3). Additionally, we found that the *cRFXa* sub-family branches closely with the animal *RFX1/2/3* sub-family (Fig 1D, Supp Fig 3) which regulates ciliogenesis across diverse animals^51, 59^. Additionally, *cRFXc* members grouped with the animal *RFX4/6/8* sub-family, whose functions are somewhat less well conserved^27, 76^. The third choanoflagellate RFX family, *cRFXb*, does not have a well-supported relationship with any specific animal RFX sub-family. We thus infer that the last common ancestor of choanoflagellates and animals expressed at least two RFX paralogs, one related to modern-day *RFX1/2/3/cRFXa* genes and the other related to *RFX4/6/8/cRFXc* genes.

### Targeted disruption of S. rosetta cRFXa delays cell proliferation and disrupts ciliogenesis

We next used CRISPR/Cas9-based gene editing^72^ to introduce an early stop codon near the 5’ ends of the *cRFXa*, *cRFXb*, *cRFXc*, and *foxJ1* genes in *S. rosetta* (Fig 2A, Supp Fig 4A, Supp File 4). Strains with truncation edits in *foxJ1*, *cRFXb*, and *cRFXc* displayed no obvious phenotypic defects (Supp Fig 4B,C). In contrast, two independently isolated *cRFXa* mutant lines, each encoding a truncated allele of *cRFXa* (*cRFXa^PTS-1^* and *cRFXa^PTS-2^*), proliferated more slowly than a wild-type control (*cRFXa^WT^*) (Fig 2B).

**Figure 2.**
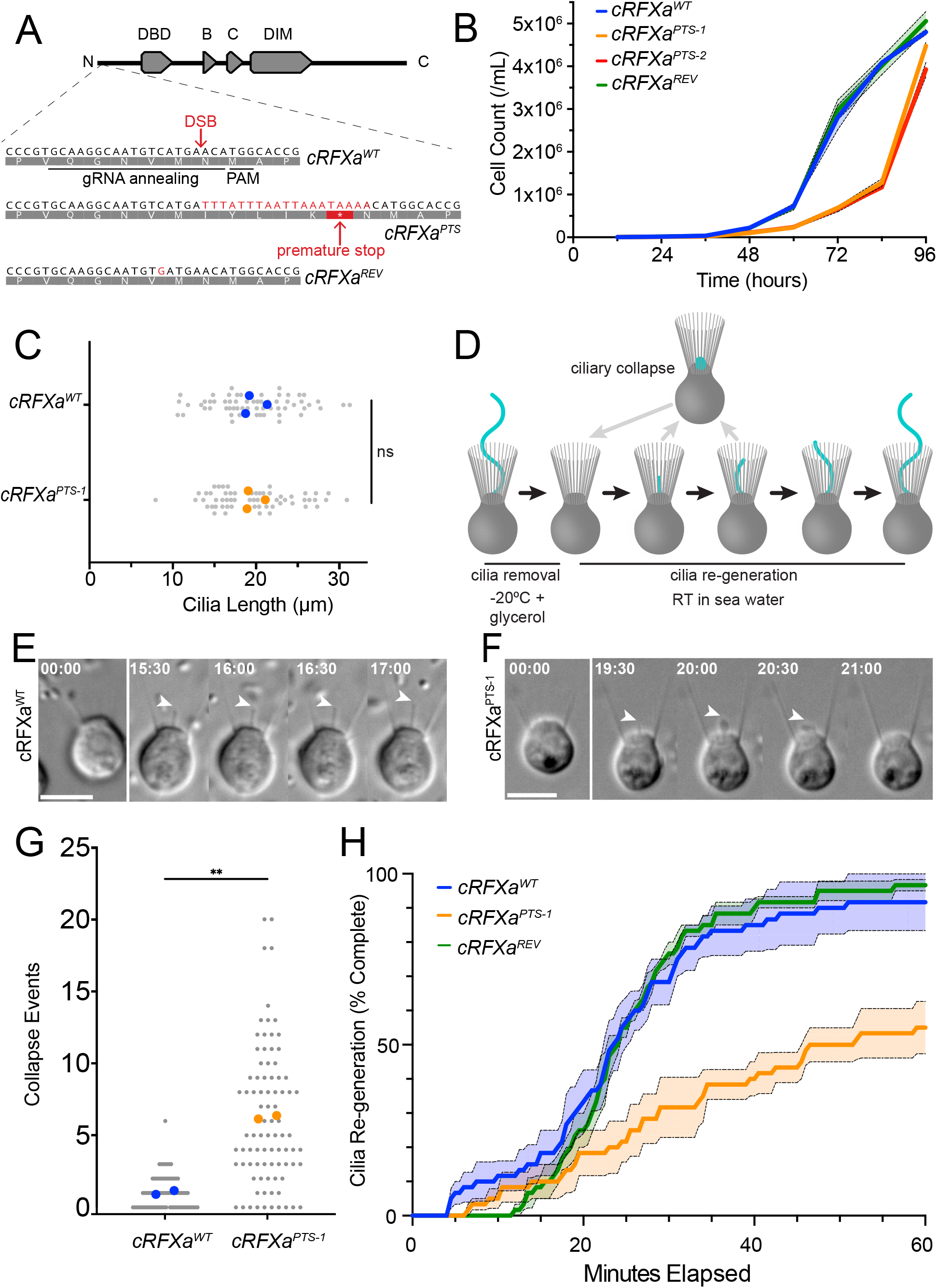
Truncation of cRFXa results in cell proliferation and ciliogenesis defects. (A) The *S. rosetta cRFXa* locus encodes a protein that contains an N-terminal DNA-binding domain (DBD) followed by two conserved domains of unknown function (B,C) and a dimerization domain (DIM)^59^. The *cRFXa* locus was targeted by a guide RNA (gRNA) that anneals in an exon near the N-terminus before the DBD, coupled with a homology-directed repair template that inserts a cassette (TTTATTAATTAAATAAA) that encodes an early stop codon (* in translation product, grey shaded letters). The edited allele is called *cRFXa^PTS^* (for Premature Termination Signal^72^) and codes for a truncated polypeptide of 24 amino acids. Two independent *cRFXa^PTS^*mutants, *cRFXa^PTS-1^* and *cRFXa^PTS-2^*, were recovered. The *cRFXa^PTS-1^* strain was reverted to a wild-type polypeptide sequence [with a synonymous GTC®GTG (Valine)], to create the *cRFXa^REV^* strain. DSB = double-strand break, PAM = protospacer adjacent motif. Numbers indicate amino acid positions in coding DNA sequence. (B) Truncation of cRFXa (*cRFXa^PTS^*) results in delayed cell proliferation compared to *cRFXa^WT^* and *cRFXa^REV^* cells. *cRFXa^PTS-1^* and *cRFXa^PTS-2^* clones were assayed. Cells were diluted to 1,000 cells/ml and triplicate samples were collected and counted every 12 hours for 96 hours. The mean values were plotted with the standard error of the mean shown as dotted lines. (C) Cilia lengths were comparable between *cRFXa^W^*^T^ (19.73 µm) and *cRFXa^PTS-1^* (19.63 µm) cells. Randomly selected cells from three biological replicates were analyzed (see Materials and Methods), measuring 20 cells per genotype per replicate, for 60 cells total per replicate. Colored dots show replicate mean values and grey dots show the lengths of individual cilia. Unpaired t-test p-value = 0.959. (D) Choanoflagellate ciliogenesis can be synchronized and quantified following ciliary removal. To this end, *S. rosetta* cells were treated with 10% glycerol at -20°C in artificial sea water, followed by a return to 100% sea water at room temperature (RT). Cilia were severed during the treatment, while the rest of the choanoflagellate cell morphology was maintained. We observed that nascent cilia can collapse and resorb before a new round of ciliary growth begins. The point at which the growing cilium passed the edge of the microvillar collar was used as a marker of successful ciliogenesis. (E) A representative time series showing a *cRFXa^WT^* cell in the process of ciliogenesis, from cilia removal (00:00 mm:ss) to growth (15:30-17:00 mm:ss). The nascent cilium (arrowhead) extends as a thin, straight protrusion; ciliary beating has not begun yet. Scale bar = 5 µm. (F) A representative time series showing a *cRFXa^PTS-1^* cell in the process of ciliogenesis. Arrowhead marks a nascent cilium that collapses (20:00 time point) and resorbs back into the cell. Resorption here is complete in one minute, which was typical. Scale bar = 5 µm. (G) Nascent cilia in *cRFXa^PTS-1^* cells collapse more frequently than *cRFXa^WT^* cells during the ciliogenesis assay. For each of two biological replicates, 20+ cells were scored for the number of ciliary collapses during a 60-minute ciliary re-growth period. Colored dots show mean values of each biological replicate and grey dots show values for individual cells. The mean number of collapses (across biological replicates) was 1.00 collapses/cell/60 minutes for *cRFXa^WT^* and 6.24 for *cRFXa^PTS-1^*. Unpaired t-test p-value = 0.0012. (H) *cRFXa^PTS-1^* cells are delayed in ciliary re-generation relative to *cRFXa^WT^* and *cRFXa^REV^* cells. Graph shows the percent of cells that have completed ciliary re-generation as a function of time (three biological replicates, 20 cells each). Regeneration was defined as the point at which the cilium grows past the collar. Dotted lines show standard error of the mean across three replicates.

To ensure that the proliferation defect of the *cRFXa^PTS^*mutants was caused by the introduced mutation rather than an undetected off-site mutation, we reverted the *cRFXa^PTS-1^* strain using a CRISPR repair template that restored the wild-type amino acid sequence. In addition to removing the early stop codon, the engineered revertants also encoded a synonymous GTC (Val) -> GTG (Val) mutation at amino acid position 18, allowing us to distinguish their genotype from that of *cRFXa^WT^* cells (Fig 2A). Growth of the *cRFXa* revertant (*cRFX^REV^*) was comparable to that of *cRFXa^WT^* cells (Fig 2B), confirming that the growth defect was a direct result of the truncation of cRFXa in the *cRFXa^PTS-1^* strain.

Because RFX1/2/3 homologs are essential for proper ciliogenesis in animals^59^, we investigated whether the *S. rosetta cRFXa^PTS-1^* mutant might display ciliary defects. Under standard growth conditions (Materials and Methods), cilia lengths were indistinguishable between populations of *cRFXa^WT^* cells (mean = 19.73 µm) and *cRFXa^PTS-1^* cells (mean = 19.67 µm) (Fig 2C). However, measuring steady-state cilia lengths did not allow us to assess the dynamics of ciliogenesis itself.

To analyze the dynamics of ciliary re-generation, we adapted a ciliary removal protocol^77^ based on cold shock treatment with 10% glycerol. This procedure resulted in the severing of cilia while mostly preserving cell viability and morphology, including the presence of the microvillar collar (Materials and Methods, Fig. 2D-F). Upon return to room-temperature conditions in sea water, the cells almost immediately began to re-grow new cilia (Fig. 2E, Supp Movie 1). In *cRFXa^WT^* cells, a nascent cilium emerged rapidly from the apical pole of the cell and proceeded to lengthen (Fig 2E, Supp Movie 1). On occasion, the nascent cilium collapsed and was resorbed into the cell, after which the cilium began growing again (Fig 2D, Supp Movie 2). In contrast, nascent cilia in *cRFXa^PTS-1^* mutant cells frequently collapsed and were resorbed (Fig 2F,G, Supp Movie 3, 4). *cRFXa^PTS-1^* cells averaged 6.24 ciliary collapse events over 60 minutes of re-generation, while *cRFXa^WT^* cells averaged 1.00 collapse events (Fig 2G, p-value = 0.0012, unpaired t-test). These collapses were characterized by a bubble-like expansion of the ciliary membrane, followed by resorption of the ciliary membrane into the cell; each collapse and membrane resorption event was typically complete within two minutes (Fig 2F, Supp Movie 3, 4). Thus, cRFXa is required for efficient ciliogenesis.

To quantify the overall rate of ciliogenesis, we established a metric by which cells were scored as having a re-generated cilium once the apical tip of the cilium grew past the apical boundary of the microvillar collar (Fig 2D). We observed that only 55% of *cRFXa^PTS-1^* mutant cells had successfully re-generated their cilium by 60 minutes after ciliary removal, whereas 90% of *cRFXa^WT^* cells and 97% of *cRFXa^REV^* cells had completed re-generation by this time (Fig 2H). A similar ciliogenesis defect was also observed in the *cRFXa^PTS-2^* mutant (Supp Fig 4D). In contrast, the *cRFXb^PTS^*, *cRFXc^PTS^*, and *foxJ1^PTS^*mutants did not display any detectable ciliogenesis defect (Supp Fig 5A,B,C). In summary, of the genetic disruptions analyzed here, only *cRFXa^PTS^* cells displayed delayed rates of cell proliferation and aberrant ciliogenesis.

### cRFXa promotes expression of ciliogenesis genes and FoxJ1

To investigate how the disruption of *cRFXa* in *S. rosetta* leads to disruption of ciliogenesis, we next investigated the transcriptional profiles of *cRFXa^WT^* and *cRFXa^PTS-1^* cells (Fig 3, Supp File 5). In animals, the ciliary phenotypes in RFX loss-of-function studies are associated with reduced expression of ciliary genes^27–29, 31, 33, 39, 78^ and we hypothesized that the same might be true in choanoflagellates. Building on previously published databases of ciliary genes^79–81^, we curated an updated list of 269 components required for proper assembly of motile cilia in humans and whose molecular function is understood in some detail (Supp File 6). From this list of 269 human genes, we found 201 *S. rosetta* genes as likely orthologs, hereafter referred to as the “HsaSro conserved ciliome” (Supp File 6, Materials and Methods). The HsaSro conserved ciliome includes genes involved in intraflagellar transport (IFT), axonemal dyneins, radial spokes, the BBSome, tubulin modifiers, the ciliary transition zone, ciliary vesicle formation, and more (Fig. 3A).

**Figure 3.**
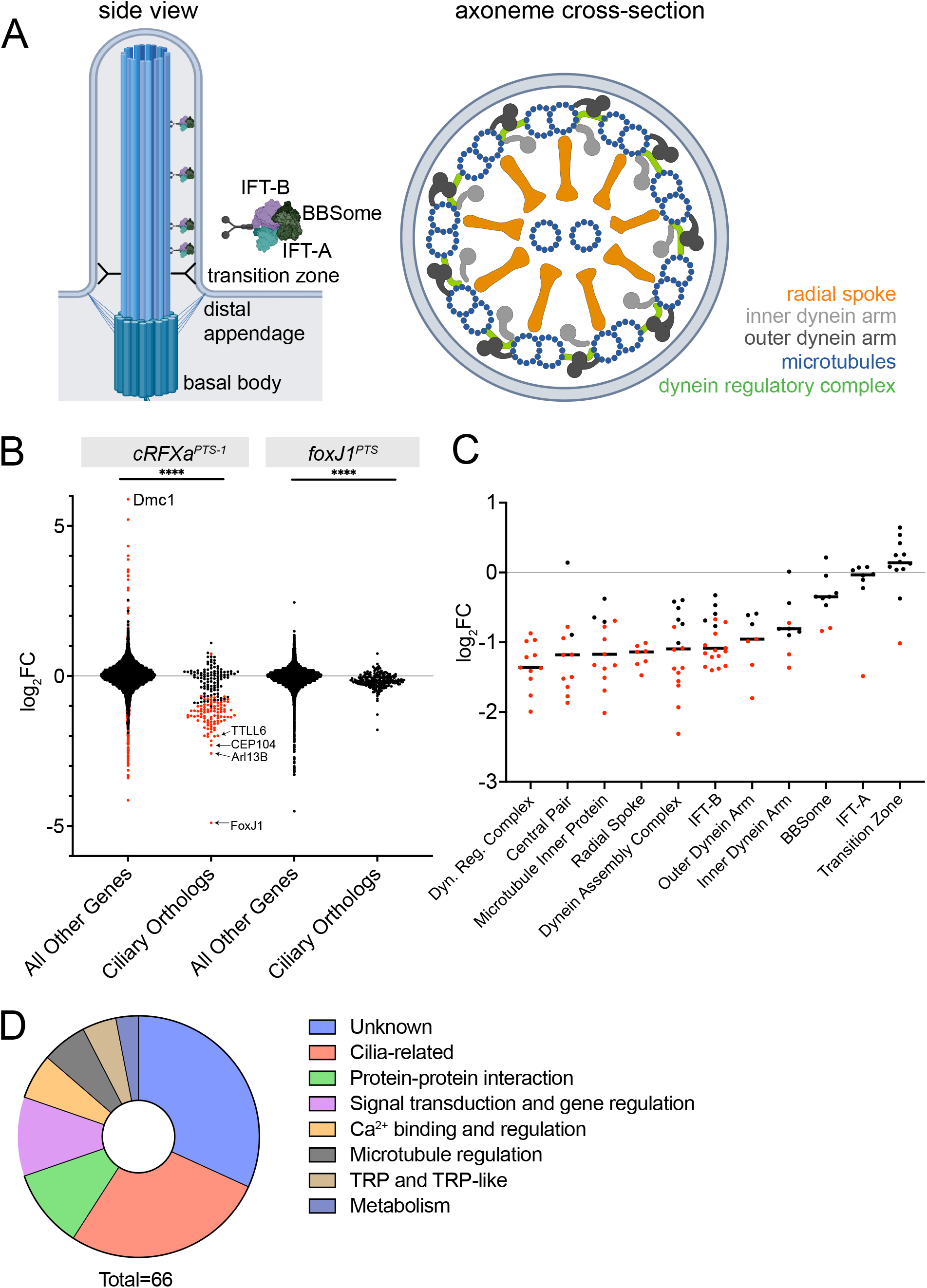
*cRFXa^PTS^* cells down-regulate conserved ciliary genes. (A) Eukaryotic motile cilia are constructed from conserved macromolecular complexes encoded by dozens of genes (Supp File 6). The side view shows how the basal body which nucleates the microtubules of the cilium docks to the cell membrane. IFT trains traverse in both anterograde and retrograde directions to shuttle ciliary components to the growing tip. Axoneme cross-section shows the organization of microtubule doublets in the cilium as well as the inter-doublet links and dynein arms that power ciliary motility. (B) Ciliary genes, including *foxJ1*, are significantly down-regulated in the *cRFXa^PTS-1^* mutant. Shown are log_2_FC values for HsaSro conserved ciliary genes (n = 201), compared to all other predicted genes in the *S. rosetta* genome. Red dots indicate genes whose differential expression was called as significant by edgeR using a false discovery rate cut-off of < 0.001. Note no individual gene was significantly differentially expressed in *foxJ1^PTS^* cells. Mann-Whitney p-value comparing ciliary orthologs to all other genes for both *cRFXa^PTS-1^* and *foxJ1^PTS^*: <0.0001. (C) Many categories of ciliary genes are down-regulated in *cRFXa^PTS-1^* cells. For each category, the horizontal bar shows the average log_2_FC value for genes in that category, while dots indicate the expression changes of individual genes. Red dots indicate a gene with an edgeR false discovery rate (FDR) < 0.001. (D) Distribution of likely functions for all genes down-regulated more than four-fold (log_2_FC < -2) in the *cRFXa^PTS-1^* mutant. Categories are based on protein domain annotation and BLAST hits.

Of the 201 genes in the HsaSro conserved ciliome, 93 were significantly down-regulated in the *cRFXa^PTS-1^* mutant (edgeR FDR < 0.001; Fig 3B, Supp File 6). The 93 down-regulated ciliary genes had slightly more than a 2-fold reduction in expression (log_2_FC = -1.34; Fig 3B), while genes not in the HsaSro conserved ciliome had on average no change in expression between *cRFXa^WT^* and *cRFXa^PTS-1^* cells (Fig 3B). Among the most down-regulated ciliary genes in *cRFXa^PTS-1^* cells were genes encoding the ciliary GTPase *ARL13B* (log_2_FC = -2.58)^82^, the ciliary tip component *CEP104* (-2.16)^83^, and the tubulin glutamylation enzyme *TTLL6* (-1.99)^84^ (Fig 3B). We confirmed that *S. rosetta* ciliary genes are preferentially down-regulated by additionally analyzing the expression pattern of all genes whose protein products were previously detected by mass spectrometry in *S. rosetta* cilia^85^ (Supp Note 3, Supp Figure 6).

Among the most down-regulated categories of ciliary genes were those involved in the dynein regulatory complex (log_2_FC = -1.33), dynein assembly factors (-1.12), radial spoke components (-1.20), and central pair components (-1.19) (Fig 3C). Interestingly, not all categories of ciliary components were affected in the mutant. Notably, genes encoding components of IFT-A and the transition zone were not down-regulated in the *cRFXa^PTS-1^* mutant (Fig 3C).

We additionally considered the functions of the most down-regulated genes in the *cRFXa^PTS-1^* mutant, regardless of whether they were members of the HsaSro conserved ciliome. Sixty-six genes were down-regulated by 4-fold or more (log_2_FC < -2) in the *cRFXa^PTS-1^* mutant (Fig 3D, Supp File 6). After manual annotation of these genes, the largest category were genes of unknown function, while the second largest category (17 genes) contains genes with a confirmed molecular role in ciliogenesis or with cilia-associated phenotypes. For example, we observed multiple genes with microtubule-binding doublecortin domains, which align most closely with the human *DCDC2* gene, which controls ciliary length, is regulated by RFX, and causes a ciliopathy^86–88^. Also observed was a relative of *NPHP3*^89^, a ciliary component and ciliopathy gene. We detected genes with likely functions in microtubule regulation, but which have not necessarily been shown to be involved in cilia biology; these may represent candidate ciliogenesis genes, given the conserved microtubule core of the ciliary axoneme. In contrast with the cell cycle regulatory function of RFX in some fungi, none of the genes in this set had clear connections to cell cycle regulation.

In addition to coordinating the gene expression of ciliary structural components, animal RFX transcription factors activate the expression of ciliary-localized membrane receptors that mediate the function of cilia as sensory hubs^39, 41, 58, 59^. These include members of the transient receptor potential (TRP) family, which are Ca^2+^ channels that can localize to cilia and be activated by mechanical, temperature, and chemical cues^3,39, 90–92.^ Calcium influences various downstream signaling pathways and modulates cilia motility in diverse eukaryotes^93–96^. In *S. rosetta*, we observed one clear TRP channel homolog and two genes with TRP-like domains down-regulated more than four-fold in the *cRFXa^PTS-1^* mutant (Fig 3D).

Among genes up-regulated in *cRFXa^PTS-1^* cells, no clear pattern presented itself. Of potential interest was the dramatic increase in transcript levels for the meiotic recombinase *Dmc1* (log_2_FC = 5.89) and the temperature-sensitive ion channel *TRPM2* (2.86), which was identified by mass spectrometry in a proteomic study of *S. rosetta* cilia^85^.

Previous work has shown that RFX and FoxJ1 cross-regulate each other’s transcription levels in animals^28, 35, 59, 62, 78^. For example*, foxJ1* is down-regulated in *Rfx3^-/-^* mouse ependymal cells, while FoxJ1 up-regulates *Rfx3* in the mouse node and is both necessary and sufficient for full *Rfx2* expression in zebrafish multi-ciliated tissues^28, 35, 62^. Intriguingly, one of the most differentially expressed genes in the *cRFXa^PTS-1^* mutant was *foxJ1*, which was 29-fold down-regulated (log_2_FC = -4.90) (Fig 3C). This raised the question of whether cRFXa regulates ciliary genes directly, or partially through the action of FoxJ1. Although no single HsaSro conserved ciliary gene was significantly down-regulated in the *foxJ1^PTS^* cells, the average expression level of the HsaSro conserved ciliome was down-regulated in *foxJ1^PTS^* compared to *foxJ1^wt^*(avg log_2_FC = – 0.04 for non-ciliome, -0.167 for ciliome) (Fig 3B, Supp File 5,6). Together with the observation that ciliogenesis proceeds normally in *foxJ1^PTS^* cells, these data suggest that under standard growth conditions *foxJ1* is a downstream target of the cRFXa regulon in *S. rosetta*, but its potential effects on ciliary gene expression are unclear.

### Predicted RFX binding sites are enriched in promoters of choanoflagellate ciliary genes

In animals, RFX directly regulates the expression of ciliary genes by binding to a GTTRCY consensus site^34, 38, 41, 42, 50, 52, 59^. RFX can bind as a monomer to one copy of this motif^53, 54^ or as a dimer to an imperfect motif palindrome called the X-box (GTNRCC N_0–3_ RGYAAC^52^), exemplified by the *H. sapiens* RFX2 motif^97^ (Fig 4A). In fungi, RFX binds to nearly identical sequences^47^, consistent with near-perfect conservation of the DNA-contacting residues in RFX DNA-binding domains^52^ (Supp Fig 1). To examine whether ciliary genes in *S. rosetta* might be directly regulated by RFX, we investigated the DNA binding preferences of cRFXa as well as motif enrichment in the promoters of *S. rosetta* ciliary genes.

**Figure 4.**
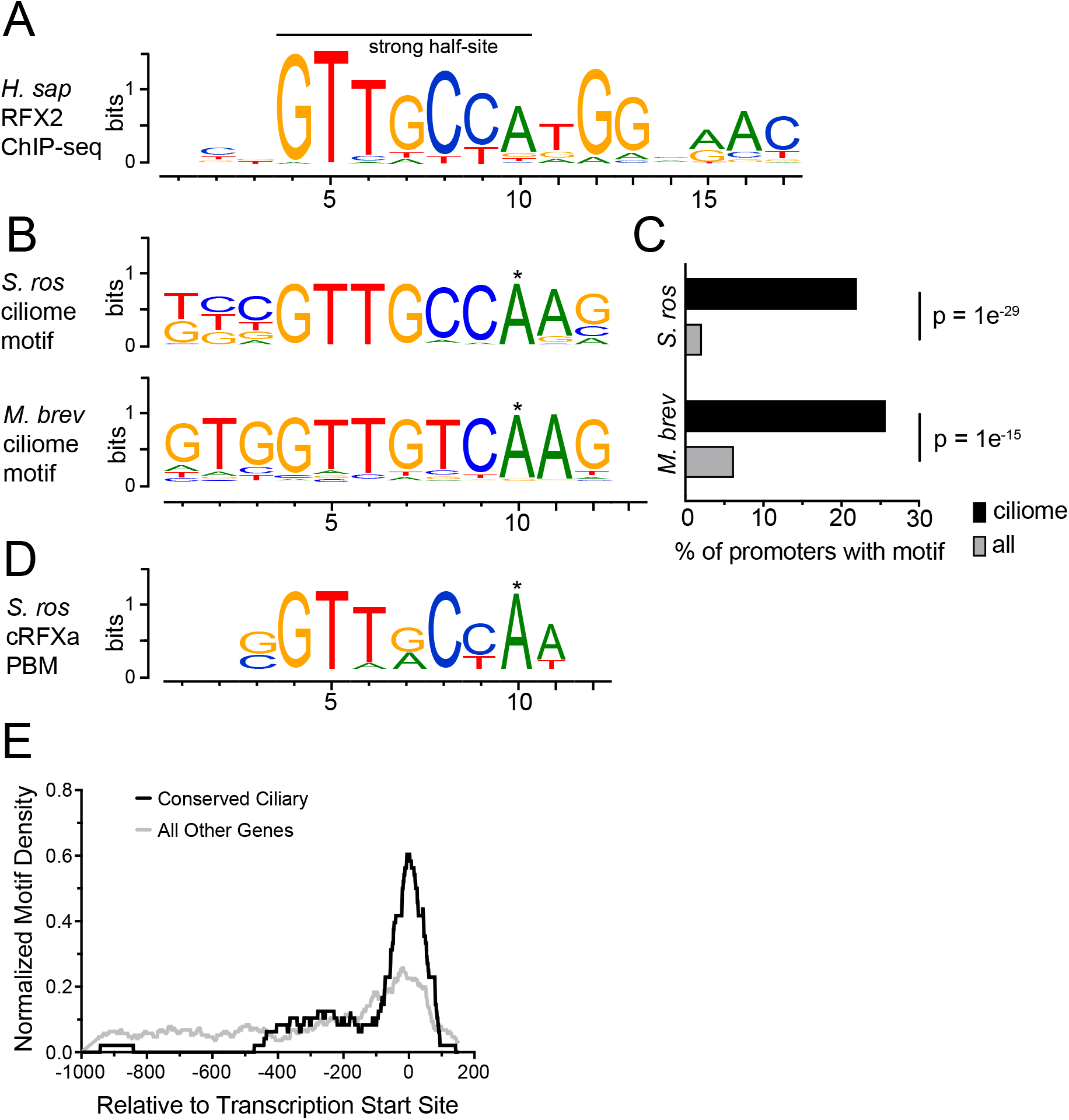
RFX motifs are enriched in choanoflagellate ciliary gene promoters. (A) The *H. sapiens* RFX2 consensus motif derived from ChIP-seq (JASPAR^97^ MA0600.1). This motif consists of two inverted, palindromic half-sites, one of which (here shown on the left) has stricter specificity requirements. The DNA binding specificity for RFX TFs is conserved across animal RFX proteins^34, 41, 52^. (B) Enriched sequence motifs in the promoters of choanoflagellate ciliary genes match RFX binding sites characterized in humans. Shown are the most enriched ciliome promoter motifs for *S. rosetta* and *M. brevicollis*, as determined by the HOMER *de novo* motif finding algorithm. Note the GTTGYCA consensus shared between the two choanoflagellate ciliome-enriched motifs and the *H. sapiens* RFX2 motif half-site. This represents the binding specificity of a single RFX DBD. For HOMER, ciliome promoters were defined as 1000 bp upstream and 200 bp downstream of annotated transcription start sites of conserved ciliome genes (Supp File 5). Asterisk indicates a position not shared by animal RFX motifs. (C) Percentage of choanoflagellate ciliome promoters with RFX-like motif compared to all mRNA promoters for both *S. rosetta* and *M. brevicollis*. RFX motifs are significantly enriched in ciliome promoters compared to all promoters, with enrichment p-values reported by HOMER. (D) The DNA binding specificity of *S. rosetta* cRFXa *in vitro*, as determined by protein binding microarray. The *in vitro* motif was built from the top ten scoring 8-mer hits (E-score range: 0.481-0.486). Asterisk indicates a position not shared by animal RFX motifs. (E) In HsaSro conserved ciliary genes, RFX motifs are preferentially located near transcription start sites. The motif density within promoters is shown for HsaSro conserved ciliome promoters and for all other promoters. The RFX motif identified by HOMER (Fig 4B) in *S. rosetta* was used. Normalized motif density (y-axis) describes the proportion of all motifs that fall into a 100 bp sliding window centered on any given position on the x-axis. The x-axis gives promoter position relative to the predicted transcription start sites of conserved ciliary genes (black line) or all other genes (grey line).

We first analyzed the promoters the HsaSro conserved ciliome (Supp File 4) using an unbiased *in silico* approach to identify any motifs that were preferentially enriched in ciliary gene promoters. Using the HOMER algorithm^98^, we detected a single motif significantly enriched in HsaSro conserved ciliome promoters. This motif showed clear similarity to published RFX binding sequences from animals (Fig 4B) and was detected in 21.9% of promoters from conserved ciliome genes (44 total) as opposed to just 2.0% of all promoters (239 total, Fig 4C). The detected enrichment of the RFX motif in HsaSro conserved ciliome promoters was robust to variable definitions of promoter length (Supp Fig 7A). In the genome of *M. brevicollis*^11^, the HOMER algorithm also detected an RFX-like motif as the most enriched motif among HsaMbrev conserved ciliome promoters (Fig 4B), with 25.6% of ciliome promoters (45 total) and 6.1% of all promoters containing occurrences of this motif (Fig 4C). We extended our analysis to *Spizellomyces punctatus,* a chytrid fungus that is ciliated and expresses RFX^99^ (Fig 1A, Supp File 1). However, our unbiased motif analysis of HsaSpunc conserved ciliome promoters did not identify any motifs (RFX or otherwise) significantly enriched.

Because the predicted choanoflagellate ciliome motifs matched functionally-validated RFX motifs from animals and fungi, we sought to confirm whether cRFXa does in fact share this binding specificity. To this end, we used an *in vitro* protein-binding microarray (PBM)^100–102^ in which full-length cRFXa from *S. rosetta* was screened against multiple panels of short DNA oligonucleotides. The consensus motif recovered by PBM (Fig 4D) showed clear similarity to both the enriched choanoflagellate ciliome motifs and the binding sites of animal RFX monomers^53, 54^ (Fig 4A). Both the computationally derived motifs from *S. rosetta* and *M. brevicollis,* as well as the PBM motif, showed an additional 3’ A (Fig 4B,D) that is not detected in animal RFX consensus sites (e.g. Fig 4A). The *in silico* detected motifs from *S. rosetta* and *M. brevicollis* promoters match the *in vitro* motif for *S. rosetta* cRFXa, and all match RFX binding sites from animals and fungi.

In animals, RFX binding motifs are enriched near transcription start sites^52, 103^. We found the same to be true in *S. rosetta*, with 60.4% of RFX-like motifs located within 50 bp of the TSS of conserved ciliary genes (Fig 4E, Supp File 7). In summary, conserved ciliary genes in choanoflagellates are enriched for predicted RFX binding sites, which are preferentially located near transcription start sites and which match the *in vitro* binding preferences of *S. rosetta* cRFXa. This additional evidence suggests that RFX directly binds to and coordinates the expression of genes required for ciliogenesis in *S. rosetta*.

## Discussion

While choanoflagellates have been reported to express RFX transcription factors, it has previously been unclear whether RFX plays a role in regulating ciliogenesis in these organisms^51, 52, 70^. Here, we provide three lines of evidence that RFX regulates ciliogenesis in the choanoflagellate *Salpingoeca rosetta*; (1) targeted disruption of cRFXa results in aberrant ciliogenesis, characterized by frequent cilia collapse events which delay the effective re-generation time after ciliary loss; (2) disruption of *cRFXa* results in wide-spread down-regulation of conserved ciliary genes, including *foxJ1*; (3) an unbiased *in silico* approach identified a conserved RFX motif as the most enriched motif in ciliome promoters compared to all promoters. The motif identified is nearly identical to the *in vitro*-derived DNA binding specificity of cRFXa, and motif occurrences cluster preferentially at transcription start sites.

In addition to the defect in ciliogenesis, disruption of *cRFXa* also results in delayed cell proliferation. Delayed cell proliferation and defective ciliogenesis may be functionally linked, as ciliary function is essential for bacterial prey capture in *S. rosetta*^104^. Post-mitotic *cRFXa^PTS^* mutant cells may experience nutrient limitation due to aberrant and delayed ciliogenesis^12^. Because choanoflagellates re-generate their cilium after mitosis^12^, it is possible that cRFXa affects non-ciliogenesis aspects of the cell cycle, as seen in fungi^45, 48^. However, our transcriptional analysis did not uncover a connection with cell cycle regulation.

cRFXa is also not entirely essential for ciliogenesis in *S. rosetta*, despite the observed defects. *cRFXa^PTS-1^* cells do eventually assemble cilia, whose lengths match those of *cRFXa^WT^*cells. This suggests a robustness to ciliogenesis, allowing cells to accommodate wide-spread down-regulation of ciliary components. It also suggests the presence of other transcriptional regulators that ameliorate the loss of cRFXa. Although cRFXb and cRFXc would be likely candidates to compensate for loss of cRFXa function, neither are appreciably expressed in *cRFXa^WT^* cells or *cRFXa^PTS-1^* (Fig 1C, Supp File 5).

The role of cRFXa in *S. rosetta* ciliogenesis, coupled with the role of the RFX1/2/3 family in animal ciliogenesis^40, 59^, and the orthology (reported here and by Chu et al^51^) between these two gene families, makes it most parsimonious that the common ancestor of animals and choanoflagellates contained an RFX transcription factor with a role in regulating ciliogenesis. This connection evolved in a step-wise fashion, as cilia were likely present in the last common ancestor of eukaryotes, while RFX transcription factors likely evolved later, in the stem lineage of opisthokonts and amoebozoans. Functional data on ciliated opisthokonts outside of the Choanozoa are missing, but our bioinformatic analysis of ciliome promoters in the chytrid *S. punctatus* did not suggest RFX involvement.

There are at least two possible interpretations for these data. In one scenario, RFX originally had a non-ciliogenesis function and was co-opted to regulate ciliary genes in the Choanozoan stem lineage. Alternatively, RFX ancestrally regulated ciliogenesis in stem opisthokonts, but was recruited for other functions in fungi, including in chytrids. In either scenario, the divergence of RFX functions between choanozoans and fungi required concerted changes in the cis-regulatory sequences of ciliary genes.

Two RFX sub-families are shared between animals and choanoflagellates: cRFXa is related to animal RFX1/2/3, while cRFXc is related to animal RFX4/6 (Fig 1D, Supp Fig 3), suggesting that an ancestral RFX gene may have duplicated before the divergence of animals and choanoflagellates. One sub-family regulates ciliogenesis in both animals (RFX1/2/3) and choanoflagellates (cRFXa), while the other shared RFX TF (RFX/4/6 and cRFXc) does not appear to regulate choanoflagellate ciliogenesis and shows mixed results for animal ciliogenesis^27, 76^. Therefore, the RFX duplication may have enabled a novel function for one RFX sub-family, namely the co-option of ciliary gene regulation. Alternatively, if the ancestral RFX already regulated ciliary genes, the duplication may have allowed the partitioning of functions between RFX sub-families.

One question raised by this work is how the ancestral RFX-ciliogenesis module was integrated into animal developmental programs. Was RFX expression sufficient for specifying ciliated cells or did it require accessory regulators? If the ancestral RFX had any non-ciliogenesis roles, how was pleiotropy resolved when re-purposing this network in animal cell type evolution? Finally, the status of FoxJ1 in these regulatory networks is intriguing. In animals, FoxJ1 operates as a master regulator of many ciliogenesis genes^35^ and shows cross-regulation with RFX. The cross-regulation of these families is also seen in *S. rosetta*, as *foxJ1* is one of the most down-regulated genes upon *cRFXa* disruption. However, disruption of *foxJ1* itself in *S. rosetta* has no detectable effect on ciliogenesis efficiency and little effect on the expression of HsaSro conserved ciliary genes in the slow swimmer cell type. This raises the question of whether FoxJ1 was a sub-module of cRFXa ancestrally and was later “promoted” to a higher level of the gene regulatory hierarchy, or whether the role of FoxJ1 in *S. rosetta* is reflects a diminishment of its role compared to its ancestral counterpart. Alternatively, disruption of *foxJ1* in *S. rosetta* may be compensated for by the activity of other TFs or the *foxJ1^PTS^* allele may not be a true knockout (e.g. alternative translation start site).

Finally, our data may add something useful to a growing discussion on the origins of animal cell types. Proposed modes and drivers of cell type evolution include division of labor^105, 106^, integration of life cycles^8, 107^, stress responses^108^, and gene, or genome, duplication^109, 110^. A common theme in many of these models is the re-purposing of ancestral regulatory connections in novel cell types, in which a single transcription factor can coordinate the activity of a suite of genes sharing complementary functionality. The work reported here provides a concrete example of a pre-animal regulatory module whose regulation was modified during the evolution of animal development to differentiate ciliated from non-ciliated cells.

## Materials and Methods

### BLAST searches for RFX and FoxJ1 genes

To determine the presence of RFX genes throughout eukaryotic diversity, we used a variety of functionally validated RFX DBDs as BLAST queries, searching against the EukProt database, which includes 993 species^73^. First, to define the broad phylogenetic distribution of RFX genes, we queried the DBDs of *Xenopus laevis* RFX2 and *Saccharomyces cerevisiae* RFX1 against the EukProt Comparative Set of 196 species, chosen for taxonomic diversity and genome/transcriptome completeness. EukProt implements the BLASTP 2.13.0 algorithm. We defined bone fide RFX hits as those with at least 75% query coverage and at least 30% sequence identity (see Supp File 1 for DBD probe sequences and EukProt BLAST results).

To develop a comprehensive set of amorphean RFX hits, we used six RFX DBD sequences (*X. laevis* RFX2, *S. cerevisiae* RFX1, *M. musculus* RFX4, *M. musculus* RFX5, *C. elegans* Daf-19, and *S. rosetta* cRFXa) as BLAST probes against a set of 95 amorphean taxa. RFX hits within these taxa were used for the data shown in Fig 1A and to construct the phylogenetic trees in Supp Fig 2 and Supp Fig 3. All sequences used for phylogenetic tree construction are detailed in Supp File 1. For *S. mediterranea*, which is of interest due to it having demonstrated FoxJ1 function in ciliogenesis, but is not hosted on EukProt, we used the BLASTP server hosted on https://planosphere.stowers.org/, which implements BLASTP 2.3.0.

We used a similar procedure to identify Fox genes, first within the EukProt Comparative Set using the DBDs from *X. laevis* FoxJ1 and *S. mediterranea* FoxJ1 as probes (see Supp File 2 for probe sequences and BLAST results) and an 75% query coverage / 30% query identity threshold criteria. To identify candidate FoxJ1 orthologs for the taxa represented in Fig 1A, reciprocal best BLAST searches were performed, using FoxJ1 DBDs from *M. musculus, X. laevis*, *S. mediterranea*, and *S. rosetta*. For these BLAST searches, we used EukProt for all except two taxa (which are not hosted on EukProt): *S. mediterranea*, hosted at https://planosphere.stowers.org/, and *X. laevis*, for which we used the NCBI BLAST server with the Uniprot reference database. In Fig1A we report taxa containing reciprocal best BLAST for either *X. laevis* or *S. mediterranea*, which are phylogenetically disparate (within animals) and both have functionally validated FoxJ1 genes with known roles in regulating motile ciliogenesis.

### Phylogenetic trees

To build maximum-likelihood trees for RFX family genes, we aligned protein sequences with MAFFT^111, 112^ (v. 7.312) using default options, trimmed with ClipKIT^113^ (v 1.3.0) using the default smart-gap trimming mode, and built trees with IQ-TREE^114^ (v. 2.2.0-beta COVID-edition) using ModelFinder^115^ and 1000 Ultrafast Bootstraps^116^. Trees were visualized with iTOL^117^. The substitution model for both the choanoflagellate RFX tree and the amorphean RFX tree was Q.pfam+F+R5.

The protein sequences used for phylogenetic reconstruction are shown in Supp File 1. Note that we do not necessarily use all the RFX genes within a given taxon, for the purposes of both clarity of presentation and the efficiency of computational bandwidth. This is especially true for vertebrates, with their abundance of RFX duplications within well-established sub-families (e.g. RFX1/2/3 genes), and for some ichthyosporeans (e.g. *C. fragrantissima*), which contain extra RFX genes with long branches that lack consistent placement in phylogenetic re-constructions. These are likely more recent lineage-restricted duplications with extensive divergence.

### RFX DNA-binding domain alignment

Selected RFX DBD sequences were aligned using MUSCLE^118^ (v. 3.8.425) with a maximum of 8 iterations, implemented in Geneious.

### Choanoflagellate culturing

Unless otherwise specified, all experiments were performed using *Salpingoeca rosetta* co-cultured with a single prey bacterial species: *Echinicola pacifica* (ATCC PRA-390, strain designation: SrEpac). Cells were grown in artificial known sea water (AKSW) supplemented with 4% cereal grass media (CGM3) and 4% sea water complete^71^. Cells are grown at 22°C and 60% humidity. For consistency, experiments were done with cells in the mid-log phase of growth, which in this media formulation occurs between 5 x 10^5^ and 3 x 10^6^ cells/ml.

### *S. rosetta* cell type RNA sequencing and analysis

Cultures were grown in triplicate for each of four *S. rosetta* cell types. Samples of slow swimmers and rosettes were prepared from cultures of 5% SWC media inoculated with 10^4^ cells/ml of *S. rosetta* feeding on *Echinicola pacifica* bacteria, and rosettes were induced with the addition of outer membrane vesicles (OMVs) from *Algoriphagus machipongonensis*^14^ . Both of those cultures were grown for 48 h at 22°C to mid-log phase. Cultures of fast swimmers were inoculated the same as slow swimmers and then grown to starvation for 3 d at 22°C, at which point we transitioned the culture to 30°C for 2.75 h to increase the population of fast swimmers. Thecate cells were prepared by inoculating the HD1 strain of *S. rosetta* – a strain that maintains a higher proportion of thecate cells while also feeding on *E. pacifica* – to 10^4^ cells/ml 10% (v/v) CGM3 and then growing for 48 h at 22°C in square plates.

For each replicate of each cell type, 5 x 10^6^ cells were processed for lysis and RNA extraction. Cells were centrifuged and washed with AKSW. Thecate cells were scraped off the plate first. Cells were resuspended in AKSW, counted, and aliquoted to 10 x 10^6^ per aliquot, then resuspended in 100 µl of lysis buffer (Booth 2018): 20 mM Tris-HCl, pH 8.0; 150 mM KCl; 5 mM MgCl_2_; 250 mM sucrose; 1 mM DTT; 10 mM digitonin; 1 mg/mL sodium heparin; 1 mM Pefabloc SC; 100 µg/mL cycloheximide; 0.5 U/µl Turbo DNase; 1 U/µl SUPERaseIN. This was incubated on ice for 10 minutes, passed ten times through a 30G needle and centrifuged at 6,000 x g for 10 minutes at 4°C. The supernatant was collected, brought to 100 µl with RNAse-free water, and RNA was purified using the RNAeasy kit from Qiagen, eluting in 30 µl of water (Cat. No. 74104).

500 ng were of RNA were used for library prep, first purified with two rounds of polyA mRNA selection with oligo-dT magnetic beads and then converted to sequencing-compatible cDNA using the KAPA mRNA HyperPrep kit (KAPA biosystems, Cat. No. KK8580), using the KAPA single-indexed adapter kit for multiplexing (KAPA biosystems, Cat. No. KK8701). RNA integrity was assessed by Agilent Bioanalyzer 2100 before library prep using an Agilent RNA 6000 Nano Kit (Cat. No. 5067-1511). Sequencing libraries were also confirmed by Bioanalyzer 2100 for the correct size distribution using the Agilent High Sensitivity DNA Kit (Cat. No. 5067-4626). Library concentration was quantified by Qubit and libraries were pooled at equal concentrations before sequencing.

Library sequencing was performed by the QB3-Berkeley Genomics core labs (QB3 Genomics, UC Berkeley, Berkeley, CA, RRID:SCR_022170). Sequencing was performed in one lane on the Illumina HiSeq 4000, collecting between 12.4 million and 61.3 million reads for each sample. Reads were de-multiplexed, checked for quality with fastqc (v 0.11.9), and aligned to predicted transcripts from the *S. rosetta* genome^119^ using Salmon^120^ (v 1.5.2.) and called for differential expression using edgeR^121^, both implemented within the Trinity software package^122^ (v 2.14.0). TPM values for RFX gene expression amongst the different cell stages, as well as differential expression tests comparing slow swimmers with thecate cells, are available in Supp File 3.

### CRISPR guide RNA and repair template design

Candidate guide RNA sequences were obtained for each gene of interest using the EuPaGDT tool (http://grna.ctegd.uga.edu/) and the *S. rosetta* genome^119^. Guide RNA length was set at 15 and an expanded PAM consensus sequence, HNNRRVGGH, was used. Coding sequences for genes of interest are easily obtained from the Ensembl Protists hosting of the *S. rosetta* genome. Guide RNA candidates were filtered for guides with one on-target hit (including making sure the guides do not span exon-exon boundaries), zero off-target hits (including against the genome of the co-cultured bacterium *E. pacifica*), lowest strength of the predicted secondary structure (assessed using the RNAfold web server: http://rna.tbi.univie.ac.at/cgi-bin/RNAWebSuite/RNAfold.cgi), and annealing near the 5’ end of the targeted gene, particularly before the region encoding the DNA-binding domain. crRNAs with the guide sequence of interest, as well as universal tracrRNAs, were ordered from IDT (Integrated DNA Technologies, Coralville, IA).

Repair templates were designed as single-stranded DNA oligos, in the same sense strand as the guide RNA, with 50 base pairs of genomic sequence on either side of the DSB cut site. Between the homology arms is the TTTATTTAATTAAATAAA insertion cassette. Repair oligos were ordered from IDT as Ultramers.

#### Genome editing

48 h prior to the transfection, *S. rosetta* cells were inoculated in 120 ml of media at 8,000 cells/mL. This seeding density brings the culture to mid-log phase at the time of transfection. Prior to the day of transfection, dried crRNA and tracrRNA from IDT were each resuspended in duplex buffer (30 mM HEPES-KOH pH 7.5; 100 mM potassium acetate, IDT Cat. No. 11-0103-01) to a concentration of 200 µM. Equal volumes of crRNA and tracrRNA were mixed, incubated for 5 minutes at 95°C in an aluminum heating block, and then cooled to 25°C slowly by removing the heat block from the heating source (with the tube still in it) and cooling to RT. The annealed crRNA/tracrRNA is referred to as the gRNA and can be stored at -20°C for weeks before use. Also prior to the day of transfection, the dried repair oligo was resuspended to 250 µM in 10 mM HEPES-KOH, pH 7.5 and incubated at 55°C for 1 hour, then stored at -20°C.

On the day of transfection, to wash away bacteria from the choanoflagellates, the culture was split into three 50 ml conical tubes and centrifuged for 5 minutes at 2000 x g. The cell pellets were resuspended and combined in 50 ml of AKSW, followed by a 5 min spin at 2200 x g. The cells were washed once more with 50 ml AKSW and spun at 2400 x g. The pellet is resuspended in 100 µl AKSW and diluted 1:100 in AKSW for counting. Cells are diluted to 5 x 10^7^ / mL in AKSW, then 100 µl aliquots (with 5 x 10^6^ cells each) are prepared.

Priming buffer is prepared by diluting 10 µl of 1 mM papain (Sigma-Aldrich Cat. No. P3125-100MG) in 90 µl of dilution buffer (50 mM HEPES-KOH, pH 7.5, 200 mM NaCl, 20% glycerol, 10 mM cysteine, filter-sterilized and stored in aliquots at -80°C). This is then diluted 1:100 in the rest of the priming buffer (40 mM HEPES-KOH, pH 7.5, 34 mM lithium citrate, 50 mM L-cysteine, 15% PEG-8000, filter-sterilized and stored in aliquots at -80°C) for a final concentration of 1 µM papain. The priming buffer can be prepared while washing the cells.

Also while washing the cells, equal volumes of pre-annealed gRNA and *Sp*Cas9 (20 µM, NEB Cat. No. M0646M) are mixed and incubated for 1 h at RT to form the RNP. 4 µl of RNP is used per transfection reaction. The resuspended repair oligo is incubated for 1 hour at 55°C to completely solubilize the material.

Each aliquot of cells is spun at 800 x g for 5 minutes and resuspended in 100 µl priming buffer and incubated for 35 minutes at RT. The priming reaction is quenched by adding 10 µl of 50 mg/ml bovine serum albumin fraction V (Thermo Fisher Scientific Cat. No. BP1600-1000). Cells are spun at 1250 x g for 5 minutes and resuspended in 25 µl Lonza SF buffer (Lonza Cat. No. V4SC-2960) if cycloheximide selection will not be used or 200 µl of SF buffer if cycloheximide selection will be used.

For each transfection, 16 µl of Lonza SF buffer is mixed with 4 µl of RNP targeting the gene of interest, 2 µl of resuspended repair oligo, and 1 µl of washed/primed cells. If cycloheximide selection is being used, 1 µl of CHX-R RNP is added as well as 0.5 µl of CHX-R repair oligo. These engineer a P56Q mutation in *rpl36a* that confers resistance to cycloheximide^72^. The nucleofection reactions are added to a 96-well nucleofection plate (Lonza Cat. No. V4SC-2960) and pulsed with a CM156 pulse in the Lonza 4D-Nucleofector (Cat. No. AAF-1003B for the core unit and AAF-1003S for the 96-well unit).

After the pulse, 100 µl of ice-cold recovery buffer (10 mM HEPES-KOH, pH 7.5; 0.9 M sorbitol; 8% [wt/vol] PEG 8000) is immediately added to each well of the nucleofection plate and incubated for 5 minutes. Then the entire contents of the well are added to 1 mL of 1.5% SWC + 1.5% CGM3 in AKSW in a 12-well plate and cultured at 22C. After one hour of culture, 10 µl of re-suspended *E. pacifica* bacteria (10 mg/ml in 1 ml AKSW) are added to each culture not undergoing cycloheximide selection, and 50 µl are added for each culture that is undergoing cycloheximide selection.

The following day, 10 µl of 1 ug/ml cycloheximide is added to wells undergoing cycloheximide selection. Selection was done for 4 days.

Clonal dilutions were done 24 hours after transfection for cells not undergoing cycloheximide selection, and 5 days after transfection (with 4 days of selection) for cells undergoing cycloheximide selection. Cells were counted and diluted to 2 cells/ml in 1.5% SWC + 1.5% CGM3 in AKSW. To this was added a 1:1000 dilution of re-suspended *E. pacifica* (10 mg/ml in 1 ml AKSW). 200 µl of diluted culture was added per well for 96-well plates. For each editing experiment, between 5 and 20 96-well plates were prepared.

To genotype, 96-well plates were screened by microscopy and wells containing choanoflagellates were marked. These were re-arrayed into fresh 96-well plates with each well containing a separate clone. To extract genomic DNA, 50 µl of cell culture was mixed with 50 µl of DNAzol direct (Molecular Research Center, Inc [MRC, Inc.], Cincinnati, OH; Cat. No. DN131), incubated at RT for 10 minutes and stored at -20°C. Genotyping PCRs were performed in 96-well plates (Brooks Life Sciences Cat. No. 4ti-0770/c) using Q5 polymerase (NEB Cat. No. M0491L), and 40 cycles of amplification. 5 µl of genomic DNA template were used in a 50 µl PCR reaction. PCR products were purified by magnetic bead clean-up and were analyzed by Sanger sequencing (UC Berkeley DNA Sequencing Facility).

### Measuring ciliary lengths

To measure cilium length, cells grown to mid-log phase were fixed and stained using 1 part Lugol’s solution (EMD Millipore Cat. No. 1.09261.1000) with 3 parts culture (usually 25 µl and 75 µl). 4 µl were loaded onto a slide, sandwiched with a No. 1.5 coverslip and imaged coverslip slide down with a Zeiss Axio Observer.Z1/7 Widefield microscope with a Hamamatsu Orca-Flash 4.0 LT CMOS Digital Camera (Hamamatsu Photonics, Hamamatsu City, Japan) and 40×/NA 1.1 LD C-Apochromatic water immersion objective. Images were acquired with 10 ms exposure and 8.0 V of light intensity, using the PH3 phase contrast ring. Ciliary lengths were traced and measured in Fiji^123^.

### Ciliogenesis assay

To monitor ciliogenesis, cells were grown to mid-log phase. Cells were counted and 6 x 10^6^ cells were centrifuged in a 15 ml falcon tube for 10 minutes at 2000 x g. The cell pellet was resuspended in 1 ml of 90% AKSW / 10% glycerol, added to a FluoroDish (World Precision Instruments Cat. No. FD35-100) and incubated for 7 minutes at -20°C. A second FluoroDish was treated with 10 seconds of corona discharge, then rinsed with 500 µl of 0.1 mg/ml poly-D-Lysine (Millipore Sigma Cat. No. P6407-5MG) in water. The dish was rinsed 3x with water and dried.

After incubation at -20°C, the cells were transferred to an Eppendorf tube and spun for 10 minutes at 4200 x g. The cell pellet was resuspended in 20 µl AKSW and transferred to the lysine-coated FluoroDish. A 22 mm circular diameter #1.5 coverslip (Electron Microscopy Sciences Cat No. 72224-01) was gently laid on top. The dish was positioned on the microscope stage and after the cells were brought into focus, the dish was flooded with 1 mL of AKSW to dislodge the coverslip while leaving the cells stuck to the surface. Cells were imaged with a Zeiss Axio Observer.Z1/7 Widefield microscope with a Hamamatsu Orca-Flash 4.0 LT CMOS Digital Camera (Hamamatsu Photonics, Hamamatsu City, Japan) and 100 × NA 1.40 Plan-Apochromatic oil immersion objective (Zeiss) using a differential interference contrast (DIC) filter. Images were acquired at 10 z-slices spanning 10 µm, with one stack acquired every 30 seconds for one hour. We used 12.2 V bulb intensity and a short exposure (5 ms) to best capture the position of the flagellum as it regrows.

Image analysis was done in Fiji, marking the time point at which the growing cilium crossed the outer threshold of the collar complex. Cells were excluded from analysis if it was impossible to determine this time point, most commonly because the cell was not oriented properly or because of other cells or bacteria in the vicinity. Cells were also excluded from analysis if they maintained their cilium at time 0 (occasionally a nub of a cilium had already started to re-generate by the time the cells were put on the microscope, so a pre-existing cilium was defined as a cilium greater than 2 µm in length). Finally, cells were excluded if the cell divided or fused with a nearby cell during the time-course, or if the cell was obviously dead; this could be diagnosed by the cell having irreversibly lost its microvillar collar and not making any attempts to re-form the cilium or collar.

### Growth curves

Cells in mid-log phase were diluted to 5,000 / ml and supplemented with 10 µg/ml *E. pacifica* bacteria (diluted 1:1000 from a stock of 10 mg/ml in AKSW). 500 µl of culture was aliquoted into each well of a 24-well plate (Fisher Scientific Cat. No. 09-761-146) and cultured at 22°C. Plates were kept in a Tupperware box with dampened paper towels and the lid loosely affixed to prevent cultures from drying out but to allow gas exchange.

Every 12 hours for 96 hours, 3 wells from each strain were fixed with 10 µl of 16% paraformaldehyde (Fisher Scientific Cat. No. 50-980-487) and stored at 4°C. After all time points were collected, each sample was counted by vortexing the sample at high speed for 10 seconds to fully mix the sample, then aliquoting 10 µl into a counting slide (Logos Biosystems Cat. No. L12001 [disposable] or L12011 [reusable]) and counting using a Luna-FL automated cell counter (Logos Biosystems, Anyang, KOR; Cat. No. L20001).

### RNA sequencing and differential expression analysis for cRFXa and FoxJ1 mutants

30 ml of cells were grown to mid-log phase. For *cRFXa^PTS-1^*, wild-type *S. rosetta* was used as the wild-type comparison strain. For *foxJ1^PTS^*, which was isolated using cycloheximide resistance selection and contains the co-edited *rpl36a^P56Q^* allele, the wild-type comparison strain was a clone with only the *rpl36a^P56Q^* mutation^72^. Three biological replicates were prepared, each on a separate day, processing one wild-type and one mutant culture at a time for cell lysis and RNA extraction.

For each replicate of each strain, 5 x 10^6^ cells were processed for lysis and RNA extraction. Cells were centrifuged and washed with AKSW. Cells were resuspended in AKSW, counted, and aliquoted to 10 x 10^6^ per aliquot, then resuspended in 100 µl of lysis buffer. This was incubated on ice for 10 minutes, passed ten times through a 30G needle and centrifuged at 6,000 x g for 10 minutes at 4°C. The supernatant was collected, brought to 100 µl with RNAse-free water, and RNA was purified using the RNAeasy kit from Qiagen, eluting in 30 µl of water (Cat. No. 74104).

Library preparation and sequencing was performed by the QB3-Berkeley Genomics core labs (QB3 Genomics, UC Berkeley, Berkeley, CA, RRID:SCR_022170). 500 ng were of RNA were used for library prep using the KAPA mRNA capture kit (Cat. No. 07962240001) for poly-A selection and the KAPA RNA HyperPrep kit (Cat. No. 08105952001). Truncated universal stub adapters were ligated to cDNA fragments, which were then extended via PCR using unique dual indexing primers into full length Illumina adapters. RNA integrity was assessed by Agilent Bioanalyzer 2100 before library prep using an Agilent RNA 6000 Nano Kit (Cat. No. 5067-1511). Sequencing libraries were also confirmed by Bioanalyzer 2100 for the correct size distribution using the Agilent High Sensitivity DNA Kit (Cat. No. 5067-4626). Library concentration was quantified by qPCR using the KAPA Library Quantification Kit (Cat. No. 079601400001) and libraries were pooled at equal concentrations before sequencing.

Sequencing was performed in one lane of an SP flow cell on the Illumina NovaSeq 6000 with an S4 flowcell, collecting between 45.4 million and 73.3 million 50 bp paired-end reads for each sample. Reads were de-multiplexed using Illumina bcl2fastq2 (v 2.20) and default settings, on a server running CentOS Linux 7. Reads checked for quality with fastqc (v 0.11.9), and aligned to predicted transcripts from the *S. rosetta* genome^119^ using Salmon^120^ (v 1.5.2.) and called for differential expression using edgeR^121^, both implemented within the Trinity software package^122^ (v 2.14.0). Transcripts with a TPM value greater than 1 for both wild-type and mutant cells were excluded from analysis. Further analysis and comparisons were done using Python scripts in Jupyter Notebook with plotting in Prism 9. TPM values for all replicates and differential expression tests are shared in Supp File 5.

### Conserved ciliome genes

Lists of evolutionarily conserved ciliary genes have been assembled by comparing datasets across eukaryotic diversity using approaches such as comparative genomics and mass spectrometry. Previous compilations of ciliary genes have been published as the Ciliary proteome database^79^, Cildb^80^ and SYSCILIA^81^.

Building on these databases, we curated our own set of human ciliary genes, focusing on components with a described functional role in ciliogenesis (Supp File 6). Our list contained 269 genes. We identified likely orthologs of these genes in *S. rosetta, M. brevicollis,* or *S. punctatus* using the criteria of reciprocal best BLAST hits or a BLAST e-value < 1e^-20^. Finally, we removed duplicate hits from a given species of interest (e.g. *S. rosetta*) to finalize a list of conserved ciliary genes, which was used for downstream analysis of RNA sequencing data and promoter motif content.

### Protein binding microarray

RNA was prepared from wild-type *S. rosetta* cells grown to mid-log phase using the methods for lysis and RNA extraction described previously (see: *S. rosetta* cell type RNA sequencing and analysis). cDNA was prepared form this RNA using the SuperScript IV reverse transcriptase kit (Thermo Fisher Scientific, Cat. No. 18091050), with 150 ng of RNA input and dT(20) primers. The cRFXa CDS was amplified from cDNA using primers MC252 and MC253 and Q5 DNA polymerase (NEB Cat. No. M0491L), with 2 µl of cDNA template in a 50 µl PCR reaction and 35x cycles. The PCR product was gel purified (Qiagen, Venlo, NLD, Cat. No. 28706) and cloned into TOPO pCR2.1 (Thermo Fisher Scientific Cat. No. K450001) after A-tailing with Taq polymerase (NEB Cat. No. M0273S) for 15 minutes at 72°C. The TOPO reaction was transformed into TOPO OneShot cells, cultured over-night, mini-prepped (Qiagen, Cat. No. 27106) and confirmed for correct insertion with Sanger sequencing (UC Berkeley DNA Sequencing Facility) using M13R primer.

The cRFXa CDS was amplified from the TOPO vector using primers MC276 and MC277 (IDT) and Q5 DNA polymerase in a 50 µl PCR reaction. The primers contain homology arms for Gibson assembly into the pTH6838 vector, which was linearized with restriction enzyme XhoI (NEB Cat. No. R016S). The pTH6838 vector is a T7-driven expression vector with a N-terminal GST tag. The amplified CDS and XhoI-digested vector were gel purified. Gibson assemblies were performed using the NEB HiFi Assembly Kit (New England Biolabs, Cat. No. E2621L) with 100 ng of insert and a 2:1 molar ratio of insert:vector. The Gibson reaction was transformed into chemically competent XL10 Gold *E. coli* (Agilent, Santa Clara, CA, Cat. No. 200315), cultured over-night, mini-prepped and confirmed for correct insertion with Sanger sequencing.

The TF samples were expressed by using a PURExpress In Vitro Protein Synthesis Kit (New England BioLabs) and analyzed in duplicate on two different PBM arrays (HK and ME) with differing probe sequences. PBM laboratory methods including data analysis followed the procedure described previously^100, 101^. PBM data were generated with motifs derived using Top10AlignZ^102^.

#### Promoter transcription factor motif analysis

From the conserved ciliary genes in *S. rosetta*, *M. brevicollis,* or *S. punctatus* (Supp File 6), we extracted the promoter regions, defined as 1000 base pairs upstream and 200 base pairs downstream of annotated transcription start sites, although other promoter definitions were tested to ascertain the robustness of the results (Supp Fig 7). Using the ciliary promoters and a background set of all promoters (-1000 to 200 bp from all protein-coding genes), we looked for ciliome-enriched motifs using HOMER^98^, specifically the findMotifs.pl script with default options. To create a list of motif instances from a HOMER-identified motif, we also called findMotifs.pl with the -find option.

For *S. rosetta*, we used gene models from assembly Proterospongia_sp_ATCC50818, hosted on Ensembl Protist. For *M. brevicollis*, we used gene models from assembly GCA_000002865.1, hosted on Ensembl Protist. For *S. punctatus*, we used gene models from assembly DAOM BR117, hosted on Ensembl Fungi.

## Supporting information

SuppFile1_RFXblast

SuppFile2_FoxBLAST

SuppFile3_life_history_expression

SuppFile4_gene_editing_information

SuppFile5_MutantRNAseq

SuppFile6_conserved_ciliome_analysis

SuppFile7_RFXmotif_instances

SuppMovie1

SuppMovie2

SuppMovie3

SuppMovie4

## Acknowledgements

We are so grateful to members of the King Lab for maintaining a fun and supportive work environment. Thanks to the following people for critical reading of the manuscript and/or valuable feedback throughout the project: Thibaut Brunet, Josean Reyes-Rivera, Michael Carver, Alain Garcia de Las Bayonas, Jacob Steenwyk, Arnau Sebé-Pedrós, Lillian Fritz-Laylin, Mike Eisen, Fyodor Urnov, Elçin Ünal, Monika Sigg, Flora Rutaganira, Erika López-Alfonzo. We thank Lily Helfrich for standardizing cell type growth conditions, Alexandra Mulligan for help with molecular cloning, Flora Rutaganira for help with screening RFX antibodies, and Jacob Steenwyk for advice on phylogenetic computational packages. We thank the UC Berkeley DNA Sequencing Facility and QB3 Genomics Facilities. MCC was supported by an NIH MCF Training Grant and an NSF Graduate Research Fellowship Program Award, and research in the King Lab is supported by an Investigator Award from the Howard Hughes Medical Institute. Work in the Hughes Lab was funded by a CIHR grant to THR (FDN-148403).

## Supplementary Material

**Supplementary Figure 1.**
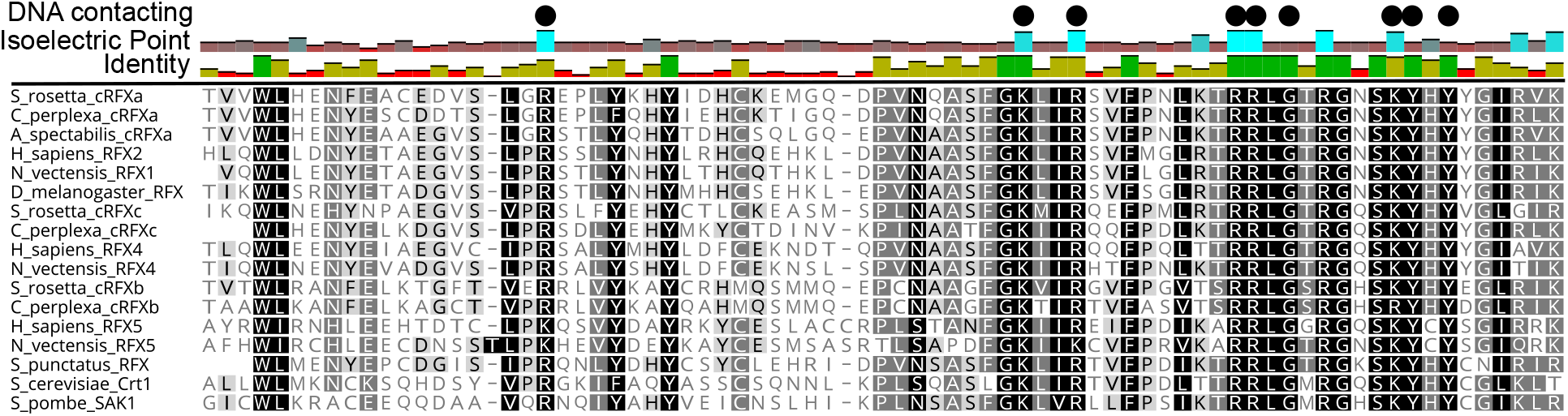
RFX DNA-binding domain sequences have highly conserved DNA contacting residues. Selected RFX DNA-binding domains were aligned with MUSCLE and individual residues shaded according to identity. DNA contacting residues as determined by a crystal structure of *H. sapiens* RFX1^50^ are labeled with black circles. These largely basic residues (note their correspondence with the average isoelectric point of each residue in the alignment) are almost perfectly conserved across all RFX sequences.

**Supplementary Figure 2.**
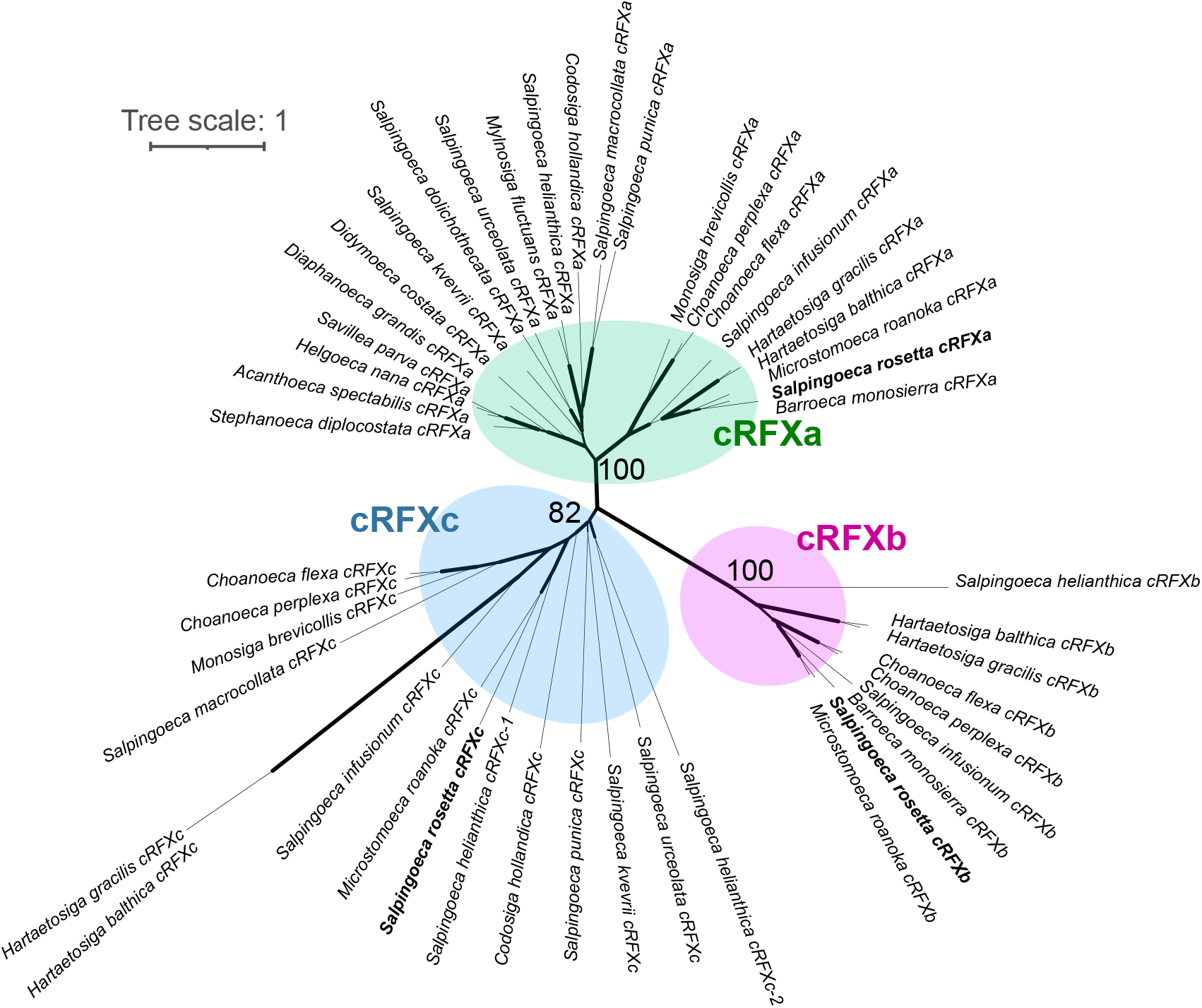
Choanoflagellate RFX genes form three well-resolved clades. Choanoflagellate RFX protein sequences were aligned with MAFFT, trimmed with ClipKIT, and assembled into a maximum-likelihood phylogenetic tree with IQ-TREE. Every choanoflagellate with RFX genes contains a copy of *cRFXa* (green, 100% UF-boot support), while *cRFXb* (pink, 100% UF-boot support) and *cRFXc* (blue, 82% UF-boot support) are found in subsets of choanoflagellate taxa. Tree scale indicates length of branch corresponding to one substitution per site in amino acid alignment.

**Supplementary Figure 3.**
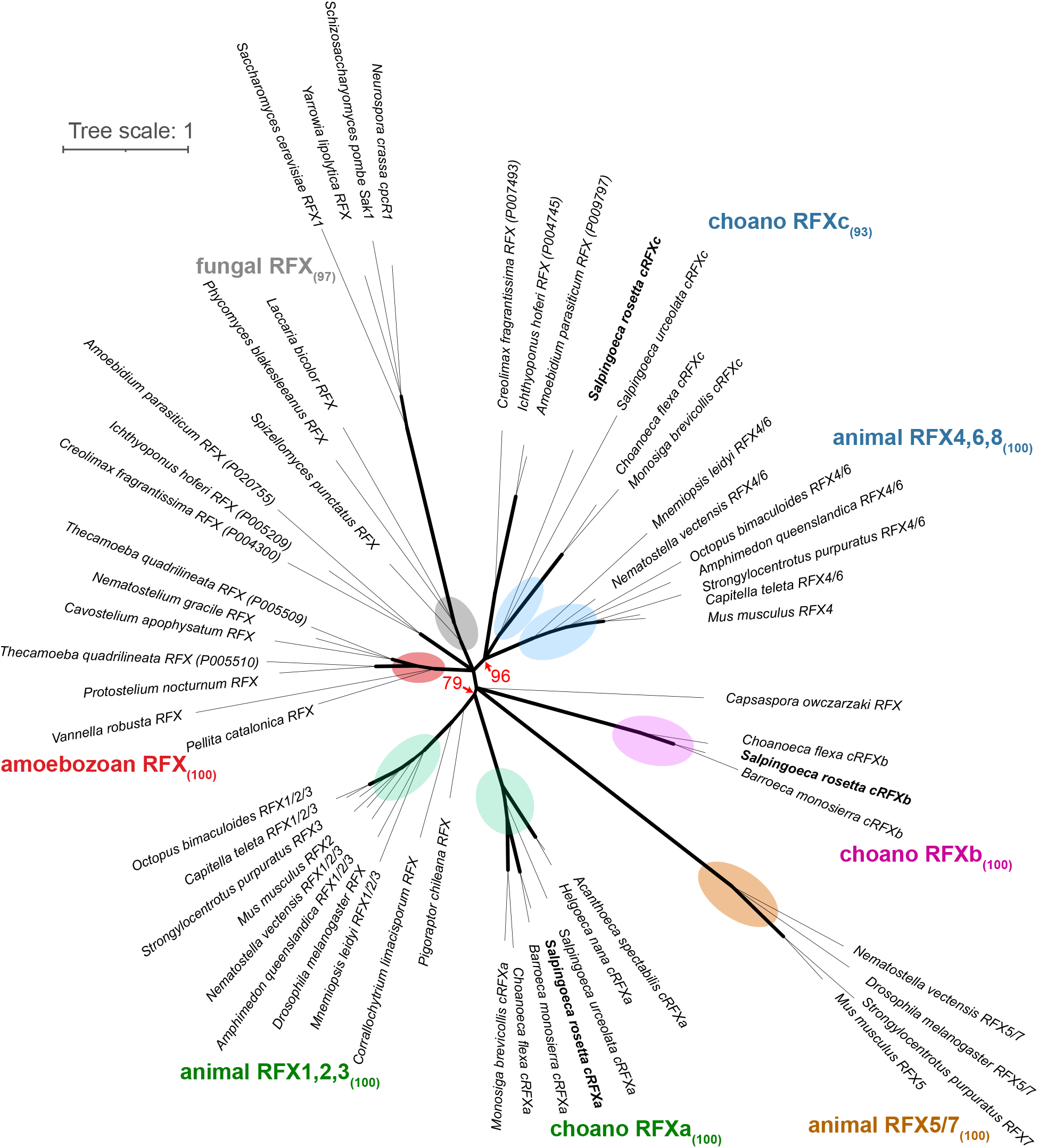
Choanoflagellate *cRFXa* genes are orthologous to the animal *RFX1/2/3* family, while *cRFXc* is homologous to the animal *RFX4/6/8* family. Selected RFX protein sequences from across diverse opisthokonts and amoebozoans (Supp File 1) were aligned with MAFFT, trimmed with ClipKIT, and assembled into a maximum-likelihood phylogenetic tree with IQ-TREE. The three previously discovered choanoflagellate RFX families were well-resolved, as were the three animal RFX families, fungal RFX genes, and amoebozoan RFX genes. Red letters and arrows indicate UF-boot support for nodes that connect animal and choanoflagellate RFX gene families. Note that ichthyosporeans (*A. parasiticum*, *C. fragrantissima*, *I. hoferi*) contain at least two RFX genes, one of which groups with *cRFXc* and *aRFX4/6/8*. Width of branches indicates bootstrap support and all nodes with less than 75% bootstrap support are collapsed. Tree scale indicates length of branch corresponding to one substitution per site in amino acid alignment.

**Supplementary Figure 4.**
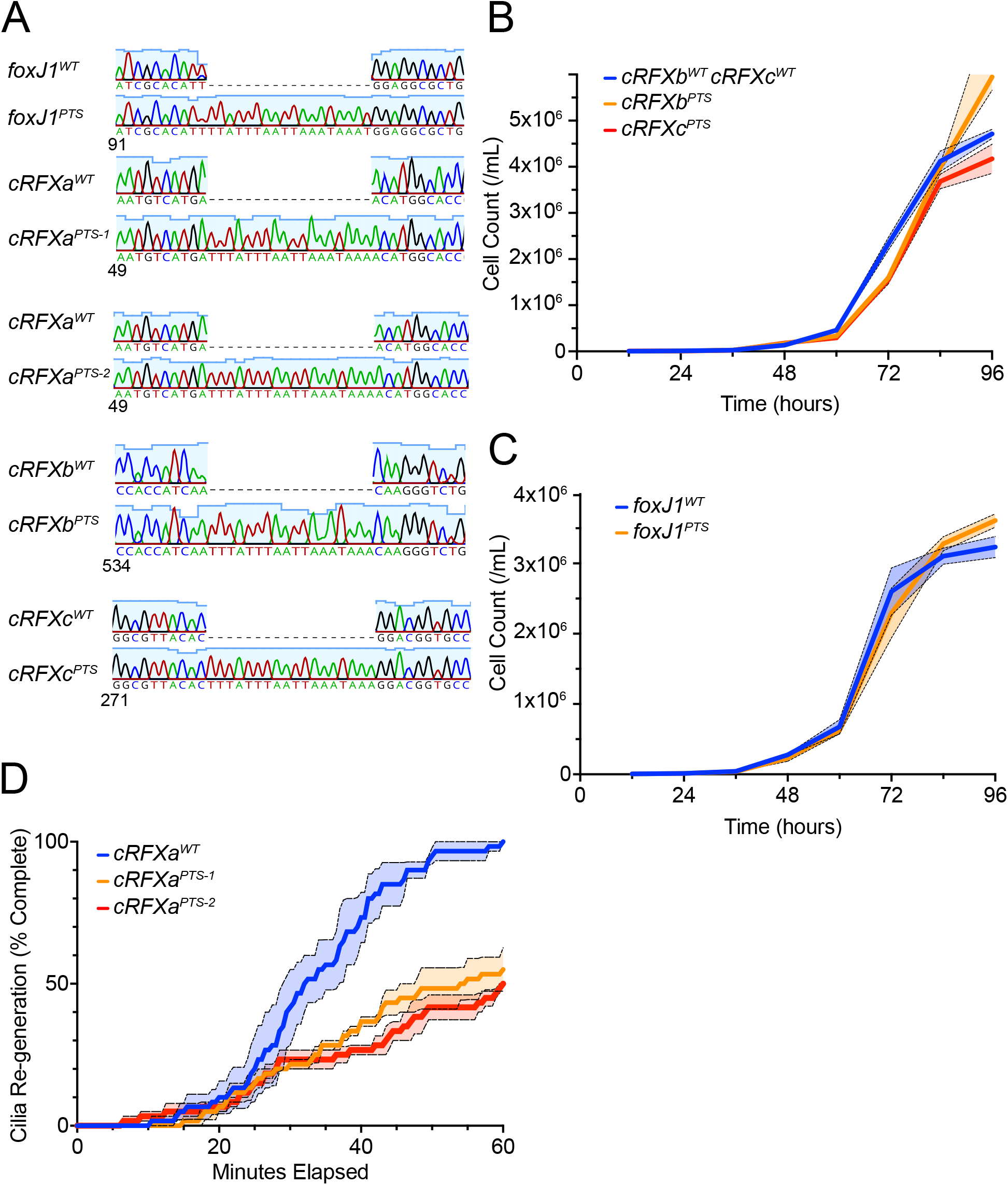
(A) Sanger sequencing confirms intended insertions at genome-edited loci. Clonally isolated cells from CRISPR genome editing experiments were genotyped by PCR and Sanger sequencing (Supp File 3). Numbers show relative position in the coding DNA sequence of the target gene. The TTTATTTAATTAAATAAA cassette is introduced in an exon and creates a stop codon in every possible reading frame. All genes are truncated before the DNA-binding domain. (B) *cRFXb^PTS^*and *cRFXc^PTS^* strains show equivalent proliferation rates compared to an isogenic strain (Materials and Methods). As in Fig 2B, cells were diluted to 1,000 cells / ml and triplicate samples were collected and counted every 12 hours for 96 hours. The mean values are plotted with the standard error of the mean shown as dotted lines. (C) *foxJ1^PTS^* shows an equivalent proliferation rate compared to an isogenic strain. Growth rates were assayed and quantified as in Fig 2B and Supp Fig 2B. (D) *cRFXa^PTS-2^*strain shows a ciliogenesis defect comparable to that observed in *cRFXa^PTS-1^*. Ciliogenesis was compared under standard growth conditions as described in Figure 2G. For each strain, triplicate experiments were done, quantifying the time point of completed re-generation for each of 20 cells, and plotting the percent that have completed ciliary re-generation as a function of time. Dotted lines show standard error of the mean across the three replicates.

**Supplementary Figure 5.**
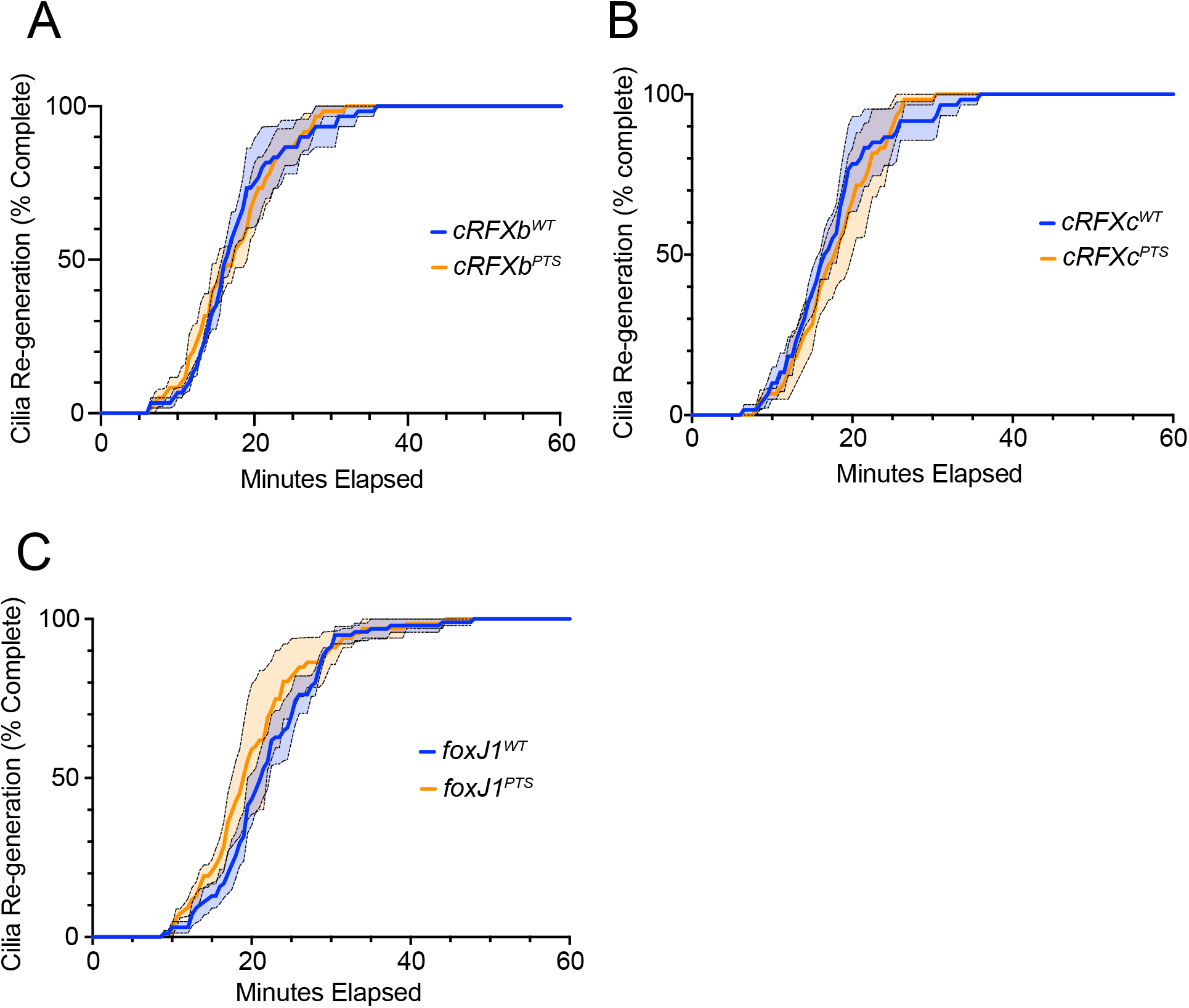
(A) The *cRFXb^PTS^*strain shows no defect in ciliogenesis. The data represents the average of three triplicate experiments (n=20 cells each) plotting the percent that have completed ciliary re-generation as a function of time. Dotted lines represent standard error of the mean (B) The *cRFXc^PTS^*strain shows no defect in ciliogenesis. The data represents the average of three triplicate experiments (n=20 cells each) plotting the percent that have completed ciliary re-generation as a function of time. Dotted lines represent standard error of the mean (C) The *foxJ1^PTS^*strain shows no defect in ciliogenesis. The data represents the average of three triplicate experiments (n=20+ cells each) plotting the percent that have completed ciliary re-generation as a function of time. Dotted lines represent standard error of the mean

**Supplementary Figure 6.**
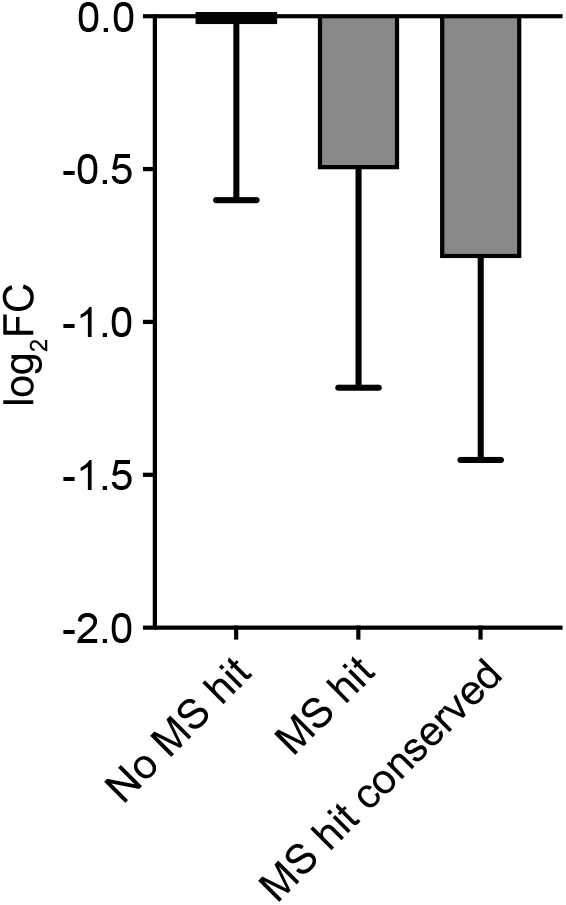
Genes with protein products identified in *S. rosetta* cilia by mass spectrometry^85^ (MS) are down-regulated in *cRFXa^PTS-1^* cells. Solid bars show the mean log_2_FC values for proteins not identified by MS in the cilia, genes that were identified by MS in cilia, and a subset of the MS hits: proteins whose sea urchin and sea anemone orthologs were also identified in the ciliary proteome of those respective taxa (MS hit conserved). Errors bars show standard deviation.

**Supplementary Figure 7.**
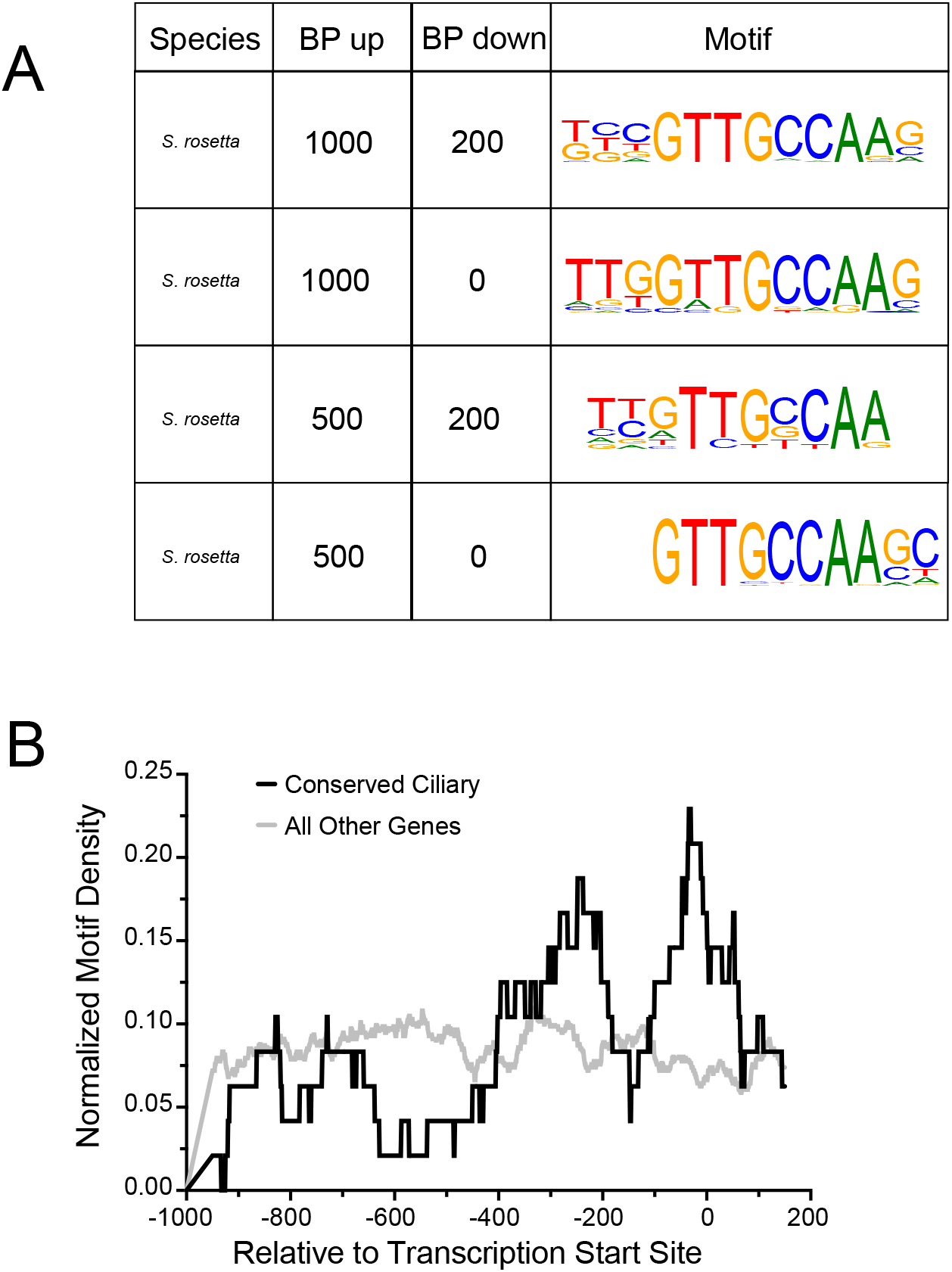
(A) RFX-like motifs are identified as the most enriched in *S. rosetta* ciliome promoters across different definitions of promoter length, relative to annotated transcription start sites. Promoters were extracted using the criteria displayed and analyzed for motif enrichment using HOMER and our set of HsaSro conserved ciliary genes (Supp File 6). (B) In ciliary genes, *M. brevicollis* RFX motifs are preferentially located near transcription start sites. The motif density within promoters is shown for motif instances in conserved ciliome genes, as well as motif instances in all other promoters. The RFX motif identified by HOMER (Fig 4B) in *M. brevicollis* was used. Normalized motif density (y-axis) describes the proportion of all motifs that fall into a 100 bp sliding window centered on any given position on the x-axis. The x-axis gives promoter position relative to the predicted transcription start sites of conserved ciliary genes (black line) or all other genes (grey line).

## Supplementary Files

1. RFX BLAST results and sequences used for phylogenetic trees
2. FoxJ1 BLAST results
3. RFX expression among choanoflagellate life history stages
4. Gene editing information
5. cRFXa and FoxJ1 differential expression data
6. Conserved ciliary gene list and RNA-seq analysis
7. Instances of RFX motifs in choanoflagellate promoters

## Supplementary Notes

1. The only surveyed choanoflagellates without a detectable RFX homolog were uncultured species whose genomes have been sequenced using single-cell technologies^129, 130^. These species show relatively lower genome completeness as measured by BUSCO^73, 131^. Therefore, the apparent absence of RFX from these species may well be artefactual.
2. When surveying the distribution of RFX and Fox genes across eukaryotic diversity, our results largely confirmed that RFX genes are widespread among opisthokonts and amoebozoans, while Fox genes are widespread among opisthokonts. However, we did observe rare exceptions to this pattern. Among 539 taxa in EukProt that are not opisthokonts or amoebozoans, three had RFX hits: *Madagascaria erythrocladioides* (a rhodophyte alga), *Gloeochaete wittrockiana* (a glaucophyte alga), and *Siedleckia nematoides* (an alveolate) (Supp File 1). Among 824 non-opisthokonts in EukProt, 14 had Fox hits (Supp File 2). For both the few RFX and Fox hits, the taxa in which they were observed were distributed across eukaryotic diversity. The only obvious pattern was that four out of the eight heterolobosean taxa hosted on EukProt contained Fox hits. Given the rare and dispersed nature of RFX and Fox hits outside of the amoebozoans/opisthokonts and opisthokonts, respectively, we interpret these hits as being more likely due to some combination of horizontal gene transfer, convergent evolution, and possibly sequencing contamination, than due to the presence of RFX or Fox genes in the last common ancestor of eukaryotes.
3. Published datasets of conserved ciliary genes do not provide an exhaustive list of ciliary components and specifically do not include taxon-specific ciliary genes. Supporting the likelihood of novel ciliary components in choanoflagellate cilia, cryo-electron tomography has revealed unique features of *S. rosetta* ciliary structure^23^. To expand our analysis, we utilized a published dataset of 464 proteins that were previously identified by mass spectrometry in the *S. rosetta* ciliome^85^. 131 of these are likely to have conserved ciliary function across Choanozoa, due to the presence of orthologs detected in the ciliary proteomes of sea urchins and sea anemones^85^. This includes the TRP-channel interacting protein Enkurin^132^, which was shown to underlie cilia-associated phenotypes in humans^85^. Transcripts whose products were detected in the ciliary proteome were on average down-regulated in cRFXa mutant cells (avg log_2_FC = -0.50; Supp Fig 6), and the subset of choanozoan-conserved ciliary genes showed more extensive down-regulation (log_2_FC = -0.79), suggesting that ciliary genes with evolutionarily conserved function have greater dependence on RFX-mediated transcriptional regulation in *S. rosetta* (Supp Fig 6).

